# Development of a three-compartment *in vitro* simulator of the Atlantic Salmon GI tract and associated microbial communities: SalmoSim

**DOI:** 10.1101/2020.10.06.327858

**Authors:** R. Kazlauskaite, B. Cheaib, C. Heys, U. Ijaz, S. Connelly, W.T. Sloan, J. Russell, L. Martinez-Rubio, J. Sweetman, A. Kitts, P. McGinnity, P. Lyons, M. Llewellyn

## Abstract

Atlantic salmon are a species of major economic importance. Intense innovation is underway to improve salmon feeds and feed additives to enhance fish performance, welfare, and the environmental sustainability of the industry. Several gut models targeted at monogastric vertebrates are now in operation. Here we report progress in the development of an Atlantic salmon *in vitro* gut model, SalmoSim, to simulate three gut compartments (stomach, pyloric caecum and mid gut) and associated microbial communities. The artificial gut model was established in a series of linked bioreactors seeded with biological material derived for adult marine phase salmon. In biological triplicate, the response of the *in vitro* system to two distinct dietary formulations (fish meal and fish meal free) was compared to a parallel *in vivo* trial over forty days. 16S rDNA sequencing, qPCR, ammoniacal nitrogen and volatile fatty acid measurements were undertaken to survey microbial community dynamics and function. SalmoSim communities were indistinguishable (p=0.230) from their founding inocula at 20 days and most abundant genera (e.g. *Psycrobacter, Staphylococcus, Pseudomonas*) proliferated the *in vitro* system. Real salmon and SalmoSim responded similarly to the introduction of the novel feed, with most taxa (96% Salmon, 97% SalmoSim) unaffected, while a subset of taxa was affected non-identically across both systems. Consistent with a low impact of the novel feed on microbial community function, VFA profiles were not significantly different in SalmoSim pre and post the switch feed. This study represents an important first-step in the development of an *in vitro* gut system as a tool for the improvement of salmon nutrition.

## Introduction

Over the last 50 years per capita fish consumption has almost doubled from 10 kilograms in the 1960s to over 19 kilograms in 2012 (FAO, 2018). The increase in the demand for fish protein has put wild fish stocks under pressure. The aquaculture sector now produces almost 50% of all fish for human consumption and has been predicted to provide 62% by 2030 (Moffitt & Cajas-Cano, 2014). The Atlantic salmon (*Salmo salar*) is the leading farmed marine fish and the 9^th^ most important aquaculture fish species farmed globally, in economic terms (FAO, 2018). Atlantic salmon are carnivores and wild pelagic fish stocks from reduction fisheries are an important protein source (fish meal (FM)), and the principle lipid source (fish oil FO), exploited to feed farmed salmon. FMFO reduction fisheries negatively impact marine ecosystems, and feeding farmed salmon on these ingredients can be unsustainable as well as expensive (Cashion et al., 2017; Worm et al., 2006). To address these issues farmed salmon feed composition has changed considerably during the relatively short history of intensive salmon farming in Norway, reducing the ratio of the marine origin components within salmon feed from around 90% in 1990 to 30% in 2013 (Ytrestøyl et al., 2015). However, there is evidence that non-marine dietary ingredient can result in the reduced fish growth rate, altered gut health as well as the modification fish gut microbial community composition and activity (Beheshti Foroutani et al., 2018; Gajardo et al., 2017a; Ingerslev et al., 2014). There is therefore considerable interest around the development of novel ingredients that have comparable performance to marine ingredient-based feeds in terms of their impact of the host and its associated microbes.

To study the impact of novel feed ingredients on gut microbial communities (e.g. Gajardo et al., 2017), as well as the addition agents (pre-biotics, pro-biotics) tailored to modify microbial community diversity and function (e.g. Gupta et al., 2019), *in vivo* trials are widely deployed in salmonid aquaculture. Although physiologically relevant, *in vivo* trials have several scientific, ethical and practical disadvantages. In salmonids, for example, gut sampling is terminal, preventing the generation of time series data from individual animals/microbial communities. Furthermore, microbial impacts on feed ingredients cannot be subtractively isolated from host enzymic/cellular activity. From an ethical perspective, *in vitro* models offer the opportunity to reduce harm via replacement of *in vivo* models (Payne et al., 2012). Practically, *in vitro* testing of salmon feed ingredients and formulations has the potential to substantially reduce the significant costs and time involved *in vivo* trials. At present, there is only one other gut system in place simulating a generalised teleost’s gut, (‘fish-gut-on-chip’ (Drieschner et al., 2019)) that exploits microfluidic technology. This system is based on the reconstruction of the rainbow trout’s intestinal barrier by culturing only intestinal cell lines in an artificial microenvironment and currently does not involve microbial communities isolated from the fish’s gut.

Prior to deploying an *in vitro* gut microbiome simulator to perform biological experiments, several criteria must be met. First, steady-state microbial communities need to be established prior to the experimental procedure to ensure that results due to experimental treatments are not confounded with bacterial adaptation to *in vitro* environment (Possemiers et al., 2004). Furthermore, physicochemical conditions within the artificial gut simulator need to be similar to those found in the gut of the target species, the bacterial communities need to be gut region-specific and representative of (if not identical to) the *in vivo* situation (Van Den Abbeele et al., 2010). Finally, the *in vitro* gut simulator needs to be validated against a parallel *in vivo* experiment, to establish to what degree the results from the experimental protocol within the artificial gut are generalisable to the *in vivo* situation (Molly et al., 1994). Several molecular techniques can be deployed to analyse microbial populations within the gut. Multiplex quantitative PCR (qPCR) in combination with taxon-specific primers can rapidly detect and quantify the bacterial groups within a large population (Postollec et al., 2011). Metagenomics sequencing approaches can provide a more detailed assessment of the microbial composition of the gut, although it may be less useful for day-to-day monitoring of specific taxa (Malla et al., 2019).

The aim of the current study is to develop a synthetic, continuous salmon gut microbial fermentation simulator, representing generalised marine lifecycle stages of Atlantic salmon. Salmonids are gastric fishes. Their guts are characterised by a clearly defined stomach followed by a pylorus with attached blind vesicles called pyloric caeca, as well as a relatively short and non-convoluted posterior (mid and distal) intestine leading to the anus (Lkka et al., 2013). Our experimental gut system simulates the stomach, the pyloric caeca and the midgut regions of the gastrointestinal tract of Atlantic salmon.

In this study, we describe the development and validation of an artificial salmon gut simulator (‘SalmoSim’) by comparing *in vivo* modulation of the gut microbial community during a feed trial with a parallel trial, performed *in vitro*. The aims of this study were to (i) develop and establish the *in vitro* system, (ii) to determine the time it takes for bacterial communities to reach steady-state conditions *in vitro* gut, (iii) analyse the similarities or differences between different the microbial communities inhabiting the different model gut compartments, (iv) evaluate similarities and differences between microbial community dynamics in the *in vitro* and *in vivo* systems.

## Materials and Methods

### Experimental set-up and sample collection in an aquaculture setting

The Atlantic salmon (*Salmo salar*) feed trial was performed by MOWI ASA at their research site in Averøy, Norway. Prior to commencement of the feed trial, salmon were fed on a fish meal diet until they reached ca. 750 grams in mass. Fish were separated into 5×5 meter marine pens (150 randomly distributed fish per pen) in 4×4 design. Four pens were randomly assigned to each of the trial diets. This study focusses on eight pens housing fish fed on fish meal and fish meal free diets (Supplementary Table 1, Figure 1F). The feed trial was conducted over five months (November 2017 - March 2018). Two randomly selected fish were collected at the end of the feed trial from the each pen assigned to the different feeds (N=6/feed) and sacrificed by MOWI employees. Stomach (N=2), pyloric caecum (N=2) and midgut (N=2) (approximately 20 cm from the vent) compartments that were collected and transferred to 15 ml Falcon tubes containing 30% glycerol and 1.5 ml cryovials and kept on ice before long term storage in −80°C freezer. Details of sample collection from farmed Atlantic salmon have been described previously (Heys et al., 2020).

**Figure 1.**
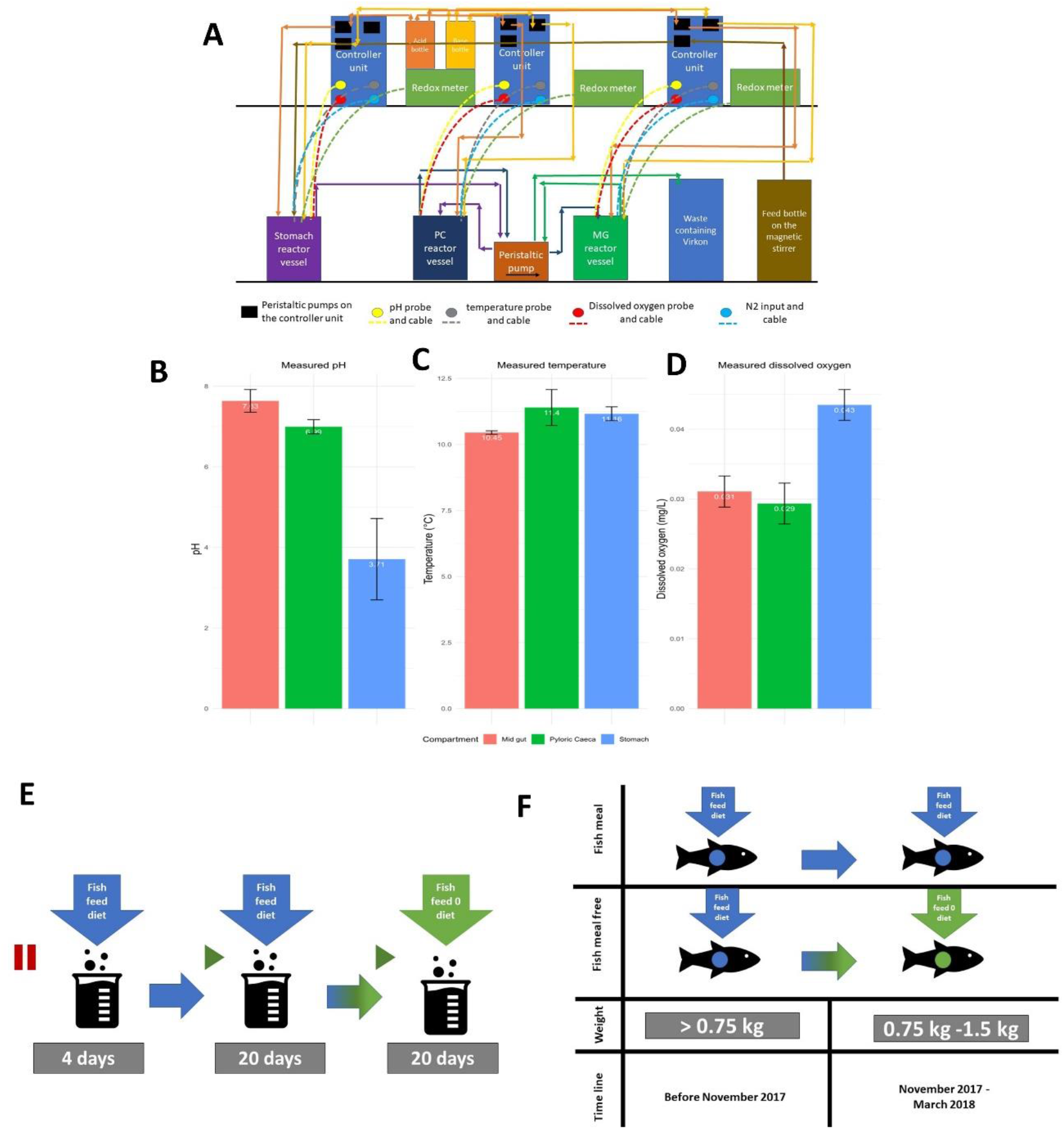
Artificial gut model system set-up, physiochemical conditions within the real salmon gut and in vivo and in vitro feed trial set up. 1A is a schematic representation of SalmoSim system; 1B-1D measured physicochemical conditions within real salmon (n=3) gut compartments: pH (1B), temperature (°C, 1C), dissolved oxygen (mg/L, 1D); 1E SalmoSim feed trial design; 1F in vivo feed trial design

### *In vitro* feed trial within SalmoSim system

#### Physiochemical conditions within Atlantic salmon gastrointestinal tract and microbiome sampling

Physicochemical conditions (temperature, pH, dissolved oxygen) were directly measured in adult Atlantic salmon from Mowi salmon farm in Loch Linnhe, Scotland (Figure 1B-D). Bacterial inoculums were prepared for the *in vitro* trial from the different gut compartments sampled from individual fish (three biological replicates, three gut compartments per fish – stomach, pyloric caecum and midgut) collected at the start if the *in vivo* feed trial in Averøy, Norway. Prior to inoculation, inoculums were stored in 15 ml falcon tubes containing 30% glycerol solution at −80°C freezer and then dissolved in 1 ml of autoclaved 35 g/L Instant Ocean^®^ Sea Salt solution. Each individual collected in Averøy formed the founder community for each replicate of the *in vitro* trial. As such each replicate of the in vitro trial represents a true biological replicate from a distinct fish.

### In vitro system ‘feed’ preparation

*In vitro* system feed media was prepared by combining the following for a total of 2 litres: 35 g/L of Instant Ocean^®^ Sea Salt, 10 g/L of the Fish meal diet or Fish meal free diet used in the MOWI feed trial (Supplementary Table 1), 1 g/L freeze-dried mucous collected from the pyloric caecum and 2 litres of deionised water. This feed was then autoclave-sterilised, followed by sieving of the bulky flocculate, and finally subjected to the second round of autoclaving.

### *In vitro* system preparation

Three 500 ml Applikon Mini Bioreactors were filled with four 1cm^3^ cubes made from aquarium sponge filters used as a surface for biofilm formation, assembled by attaching appropriate tubing and probes (redox, temperature, and dissolved oxygen), and autoclaved. Bioreactor preparation was followed by attachment of reactor vessels to the Applikon electronic control module, connection of feed, acid and base bottles (0.01M hydrochloric acid and 0.01M sodium hydroxide solutions filtered through 0.22μm polyethersulfone membrane filter unit (Millipore, USA)). Nitrogen gas was periodically bubbled through each vessel to maintain anaerobic conditions. The reactors were then allowed to fill with 400 ml of feed media. Once the system was set up, media transfer, gas flow and acid/base addition have occurred for 24 hours axenically in order to stabilise the temperature, pH, and oxygen concentration with respect to levels measured from adult salmon.

### Initial pre-growth period during *in vitro* trial

In order to allow bacterial communities to proliferate in the *in vitro* environment without washing-out, the microbial populations within the inoculum from real salmon were pre-grown inside the SalmoSim system for four days. During this phase, the system was filled with Fish meal media preparation and inoculum, and no media transfer occurred.

### Performing feed trial within SalmoSim system

After the initial pre-growth period, each validation experiment was run for 20 days while supplying SalmoSim system with Fish meal diet. After the 20 days, SalmoSim was run for 20 more days while supplying Fish meal free food. During the full 44-day experiment (4-day pre-growth period, 20-day system fed on Fish meal diet, and 20-day system fed on Fish meal 0 diet) physiochemical conditions within three SalmoSim gut compartments were kept similar to the values measured in real salmon: temperature inside the reactor vessels was maintained at 12°C, dissolved oxygen contents were kept at 0% by daily flushing with N_2_ gas for 20 minutes, and pH was kept stable in each bioreactor by the addition of 0.01M NaOH and 0.01M HCl (stomach pH 4.0, pyloric caecum pH 7.0, and mid intestine pH 7.6). During this experiment (apart from the pre-growth period) transfer rate of slurry between reactor vessels was 238 ml per day. Finally, every day 1 ml of filtered salmon bile and 0.5 ml of sterile 5% mucous solution were added to the reactor simulating pyloric caecum compartment. The schematic representation of SalmoSim system is visualised in Figure 1 A and full feed trial within SalmoSim is visually summarised in Figure 1 E.

Sampling was performed in several steps. First, samples from initial inoculums from the real salmon were collected. Once SalmoSim main experiment was started, the sampling from each bioreactor vessel was performed every second day throughout the 40-day run period (20 samplings in total). The SalmoSim samplings were achieved by collecting 30 ml of the bioreactor contents into 50 ml falcon tube, centrifuging them for 10 minutes at 5000 rpm speed, and freezing the pellets in −20°C freezer.

### Measuring nitrogen metabolism within the SalmoSim system

At each sampling point, the bacterial community activity was assessed by measuring the protein concentration using Thermo Scientific™ Pierce™ BCA Protein Assay Kit (Thermo Fisher Scientific, USA) and the ammonia concentration using Sigma-Aldrich^®^ Ammonia Assay Kit (Sigma-Aldrich, USA). Both methods were performed according to manufacturer protocol by using The Jenway 6305 UV/Visible Spectrophotometer (Jenway, USA).

### Measuring bacterial population dynamics in SalmoSim

#### Genomic DNA extraction

The DNA extraction protocol was previously described (Heys et al., 2020). In short, samples were subjected to a bead-beating step for 60 seconds by combining samples with MP Biomedicals™ 1/4” CERAMIC SPHERE (Thermo Fisher Scientific, USA) and Lysing Matrix A Bulk (MP Biomedicals, USA). Later, DNA was extracted by using the QIAamp^®^ DNA Stool kit (Qiagen, Valencia, CA, USA) according to the manufacturer’s protocol (Claassen et al., 2013).

#### qPCR analysis

The concentration of each DNA sample was measured by using Qubit^®^ fluorometer (Thermo Fisher Scientific, USA), and the dilutions were performed by using Microbial DNA-Free Water (Qiagen, Valencia, CA, USA) Inoculums from all three real salmon gut compartments were diluted to 0.25 ng/μl. SalmoSim stomach samples were also diluted to 0.25 ng/μl, and pyloric caecum and midgut SalmoSim samples were diluted to 1 ng/μl. After, the qPCR analysis was performed on each DNA sample in duplicates by using SensiFAST™ SYBR^®^ No-ROX Kit (Bioline, UK) and primer sets summarised in table 2 at a final concentration of 1 pM of each primer. Reaction conditions for all PCR reactions were 95°C for three minutes, followed by 40 cycles at 95°C for 5 seconds, 60°C for 10 seconds and 72°C for 20 seconds, followed by a final elongation step of 95°C for 10 minutes.

In order to measure the relative abundance of the target group (target determined by the specificity of the qPCR primer pairs); several steps were undertaken by adapting △△Cq method. First, the average Cq value of each primer set negative control was found. This was followed by subtraction Cq value generated by using one of the primer pairs in Supplementary Table 2, from corresponding average Cq value of the corresponding negative control (generated with the same primer pair) in order to generate value X. After, the Cq value generated by using the general primer set was subtracted from the average Cq value of the corresponding negative control (generated using general primer set) in order to generate the value Y. Finally, the value X was divided by value Y in order to find out the relative abundance of the target group with respect to the total number of bacterial 16S copies in the sample. The equations used for all these calculations are summarised below:

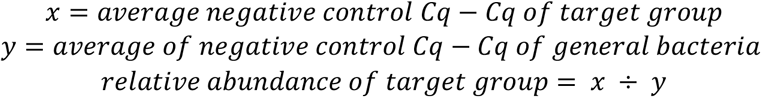

This method was carried out for each sample quantified by using different primer bacterial taxon groups. Several published and validated primer sets in the literature were used (Supplementary Table 2). Primer sets targeting *Mycoplasma*, and *Lactobacillus* genus were designed by using DECIPHER software based on the data collected by Heys et al., 2020. These primers target specificity was analysed via amplicon sequencing of the products (See Supplementary Figure 1).

#### NGS library preparation

Even though 16s ribosomal hypervariable region 4 is preferred target as it is widely used to profile vertebrate-associated microbiota, primers used to amplify this region were shown to cross-amplify *Salmo salar* 12s ribosomal gene (Heys et al., 2020; Werner et al., 2012). Amplification of the 16s V1 hypervariable region was adopted as an alternate approach (Gajardo et al., 2016). Amplification from diluted DNA samples was achieved using redundant tagged barcode 27F and 338R at final concentration of 1 pM of each primer. Primer sequences are is summarised in Supplementary Table 3. First-round PCR was performed in triplicate on each sample and reaction conditions were 95°C for ten minutes, followed by 25 cycles at 95°C for 30 seconds, 55°C for 30 seconds and 72°C for 30 seconds, followed by a final elongation step of 72°C for 10 minutes. After the triplicate reactions were pulled together into one, their concentration was measured by using Qubit^®^ (Thermo Fisher Scientific, USA), and all of them were diluted to 5ng/μl by using Microbial DNA-Free Water (Qiagen, Valencia, CA, USA). The second-round PCR, which enabled the addition of the external multiplex identifiers (barcodes), involved only six cycles and otherwise identical reaction conditions to the first. The detailed composition of second-round PCR primers is summarised in Supplementary Table 4. This was followed by the DNA clean-up by using Agencourt AMPure XP beads (Beckman Coulter, USA) according to the manufacturers’ protocol. The cleaned-up DNA was the gel-purified by using the QIAquick Gel Extraction Kit (Qiagen, Valencia, CA, USA) and then quantified by using Qubit^®^ (Thermo Fisher Scientific, USA). All the PCR products were pulled together at 10nM concentration and send for the HiSeq 2500 sequencing.

#### Statistical analysis of microbial taxon-specific qPCR data

In order to investigate the time it takes for the measured values (qPCR of different bacterial groups, protein and ammonia concentrations) to stabilise within different gut compartments of the SalmoSim system, the data for all three SalmoSim runs (three biological replicates) was combined and then sub-divided by each separate SalmoSim compartment (stomach, pyloric caecum and midgut). This data set was then further subdivided into two parts: pre- and post-feed changes. These subdivided datasets were then subjected to statistical analysis using linear mixed effect models (See Supplementary methods 1.1) to establish the effect of time on the abundance of each target taxon.

Comparisons were made between individual SalmoSim gut compartments, dataset for all three biological triplicate was combined and then subjected to statistical analysis using linear mixed effect models (See Supplementary methods 1.2) to establish the effect of gut compartment on the abundance of each target taxon.

In order to investigate if a change in the feed from Fish meal diet to Fish meal free diet results in similar trends measured in-between SalmoSim and real salmon samples, a combined data set was produced containing qPCR values measured in real salmon gut compartments (stomach, pyloric caecum and midgut of three fish fed on Fish meal diet and three fish fed on Fish meal free diet) and SalmoSim compartments at the last three time points for both feeds (once bacterial communities are stabilised while feeding SalmoSim both Fish meal and Fish meal free diets): days 16, 18, and 20 for Fish meal feed, and days 36, 38, and 40 for Fish meal free feed. This combined dataset was then separated by different SalmoSim gut compartments (stomach, pyloric caecum and midgut) and subjected to statistical analysis using linear mixed effect models (See Supplementary methods 1.3).

#### Bio-informatic analysis of 16S rRNA gene amplicon sequencing data

Sequence analysis was performed with our bioinformatic pipeline as described previously (Heys et al., 2020). First, quality filtering and trimming (>Q30 quality score) was performed on all the reads of the 16s rRNA V1 hypervariable region by using Sickle version V1.2 software (Joshi & Fass, 2011). Second, read error correction was performed by using BayesHammer module within SPAdes V2.5.0 software to obtain high-quality assemblies (Nikolenko et al., 2013). Third, paired-end reads were merged (overlap length 50bp) by using PANDAseq v2.11 software with simple_bayesian read merging algorithm (Masella et al., 2012; Schirmer et al., 2016). After overlapping, paired-end reads merged reads were dereplicated, sorted, and chimaeras (denovo and reference defined using GOLD (Mukherjee et al., 2019) and singletons were removed by using VSEARCH version 2.3.4 tool (Rognes et al., 2016). Overlapped reads were clustered in operational taxonomic units (OTUs) using VSEARCH software at 97% identity followed by a decontamination step from the host (*Salmo salar* reference genome) DNA by using DeconSeq v0.4.3 tool (Schmieder & Edwards, 2011). Taxonomic assignment of OTUs was achieved using the Scikit-learn algorithm (Pedregosa et al., 2011) implemented in QIIME 2 software and SILVA 132 database (Bolyen et al., 2019; Quast et al., 2013). Phylogenetic trees across OTUs were generated using QIIME 2 software and MAFFT for multiple sequence alignment (Katoh & Standley, 2013). OTU table was converted to a biological observation matrix (BIOM) format in order to predict the functional abundances using PICRUSt2 software (Douglas et al., 2019).

All OTU analysis was performed by using RStudio v 1.3.959 (Rstudio Team, 2019).

Alpha diversity analysis was performed by using tools supplied by Rhea pipeline (Lagkouvardos et al., 2017), microbiomeSeq package based on phyloseq package was used for ANOVA and visualisation steps (McMurdie & Holmes, 2013; Ssekagiri et al., 2017). BIOM generated OTU table was used as an input to calculate alpha diversity metrics at OTU level. Two alpha diversity metrics were calculated: microbial richness (number of observed OTUs) and Shannon diversity (considers both the number of OTUs present and their abundance per sample). Before calculating effective microbial richness, proportional filtering was performed at a relative abundance of 0.25% in each community in order to minimise the inflation in microbial richness by low abundant OTUs. In order to be able to compare Shannon index, its’ effective numbers were calculated (Jost, 2006, 2007). After, a one-way ANOVA of diversity measured between selected groups was calculated with the p-value threshold for significance set to 0.05 and its significance annotated on the plots representing different measured alpha diversity metrics.

#### Beta diversity analysis 16S sequence data

In order to investigate the effect of time on the bacterial community stability, the full dataset was used to perform beta diversity analysis using different phylogenetical distances metrics to assay community similarities between samples (weighted, balanced and unweighted UniFrac). To compare communities isolated from difference sources (SalmoSim, inoculum and real salmon), we examined a proportion of the data that contained: real salmon samples fed on Fish meal diet, all inoculum samples and stable SalmoSim time points fed on Fish meal diet (days 36, 38, and 40). This dataset was then subdivided into several different datasets with the aim of minimising the impact of rare OTUs on comparisons see details in Table 1). To investigate microbial composition differences between the gut compartments of real salmon and SalmoSim an equal number of samples (n=18) for each dataset were selected: real salmon (samples from the 3 gut compartments, from 3 biological replicates, for each of the 2 feeds) and SalmoSim (samples from each of the 3 gut compartments, for 3 biological replicate runs, for each of the 2 feeds (time point 20 for Fish meal and time point 40 for Fish meal free diet)). Finally, to establish the effect of different feeds (see supplementary Table 1 for feed formulation) on the microbial populations, three different datasets were used to perform beta diversity analysis: a dataset containing only samples from real salmon stored in −80°C freezer without glycerol; a dataset containing only samples from SalmoSim system (all data points); and a dataset containing samples only from SalmoSim once it had achieved stability (days 16, 18 and 20 for Fish meal, and days 36, 38, and 40 for Fish meal free diets).

**Table 1.** Details on the sub-setting full dataset by core OTUs. The table summarises the number OTUs within each subset dataset (subset by the % of samples that share OTUs) and percentage of the total number of OTUs within the full dataset (100%). This table also shows the number of reads and the percentage of total reads (100%) within each of the subset datasets. Note: in the 60% subset three samples were lost as they did not retain any under that criteria OTUs: Id-val1-PC1, Id-Val2-MG4 and Id-Val2-PC1.

| **% of samples that share OTUs** | **Number of OTUs (% of total OTUs)** | **Number of reads** | **% of total reads** |
| --- | --- | --- | --- |
| **80%** | 1 (0.10%) | 77,528 | 2.18% |
| **75%** | 1 (0.10%) | 77,528 | 2.18% |
| **60%** | 6 (0.61%) | 186,285 | 5.23% |
| **50%** | 13 (1.33%) | 1,179,477 | 33.10% |
| **40%** | 34 (3.48%) | 2,796,009 | 78.47% |
| **30%** | 65 (6.65%) | 3,204,411 | 89.93% |
| **100%** | 978 (100%) | 3,563,318 | 100% |

Datatsets and subsets were then used to compute ecological distances by using Bray-Curtis and Jaccards method by using vegdist() function from the vegan v2.4-2 package (Oksanen et al., 2013). Furthermore, phylogenetic distances were computed for each dataset by using GUniFrac() function (generalised UniFrac) from the Rhea package at the 0% (unweighted), 50% (balanced) and 100% (weighted) weight on the abundant lineages within the phylogenetic tree (Lagkouvardos et al., 2017). These both ecological and phylogenetical distances were then visualised in two dimensions by Multi-Dimensional Scaling (MDS) and non-metric MDS (NMDS) (Anderson, 2001). Finally, a permutational multivariate analysis of variance (PERMANOVA) by using calculated ecological and phylogenetic distances was performed to determine if the separation of selected groups is significant as a whole and in pairs (Anderson, 2001).

To provide an overall visualisation of microbial composition across all samples, a principle Coordinates Analysis (PCoA) was performed by using microbiomeSeq package based on phyloseq package (Love et al., 2017; Ssekagiri et al., 2017) with Bray-Curtis dissimilarity measures calculated by using the vegdist() function from the vegan v2.4-2 package (Oksanen et al., 2013). Bray-Curtis distances were calculated for four different datasets: the full dataset (real salmon, inoculum and all SalmoSim samples), and, three different subsets each containing only one of the free biological replicate samples from SalmoSim (Fish 1, 2, or 3), along with all real salmon and inoculum samples.

Differential abundance was calculated by using microbiomeSeq based on DESeq2 package (Love et al., 2017; Ssekagiri et al., 2017). BIOM generated OTU table was used as an input to calculate differentially abundant OTUs between selected groups based on the Negative Binomial (Gamma-Poisson) distribution.

Pearson correlation coefficient was calculated between taxonomic variables (OTUs) and two different datasets of meta-variables: (i) ammonia and protein concentrations and (ii) measured VFA values. All these correlations were calculated and visualised by using tools supplied by Rhea pipeline (Lagkouvardos et al., 2017).

#### Volatile Fatty Acid (VFA) analysis

Two stable time points for each diet were selected from SalmoSim system (for all three gut compartments) for VFA analysis: time points 18 and 20 for Fish meal diet, and time points 38 and 40 for Fish meal free diet. During runs, 1ml of supernatant from SalmoSim bioreactors was frozen in - 80°C, which was then used for VFA extraction. The protocol involved combining 1ml of supernatant with 400μl of sterile Phosphate-buffered saline (PBS) solution (Sigma Aldrich, USA) and vortexing the mixture for 1 minutes. The sample was then centrifuged at 16,000 g for 30 minutes, followed by two rounds of supernatant removal and centrifuging at 16,000 g for 30 minutes. Finally, the supernatant was then filtered through 0.2μm Costar SpinX centrifuge tube filters (Corning, USA) at 15,000 g for 2 minutes until clear. The extracted VFAs were sent for gas chromatographic analysis at the MS-Omics (Denmark).

In order to measure if the VFA concentrations are statistically different between SalmoSim fed on Fish meal and Fish meal free diet, measured VFA values dataset were subjected to statistical analysis using linear mixed effect models (See Supplementary methods 2).

## Results

### Bacterial dynamics within SalmoSim system over time

In order to explore the impact of the transfer of inoculum into the SalmoSim system on bacterial communities, as well as any potential subsequent stabilisation of these communities, alpha diversity analysis was performed on the NovaSeq output. Effective richness (Figure 2 A) estimates indicated that within the stomach and midgut compartments the initial inoculum contained the highest number of OTUs compared to later sampling time points from SalmoSim system: in stomach compartment, effective richness was statistically different between time point 0 (initial inoculum) and time points 16, 30, 36 and 38, and within midgut compartment number of OTUs within initial inoculum (time point 0) was statistically different from time points 2, 4, 6, 16, 34, 36, 38, and 40. However, within the pyloric caeca compartment, only one-time point (time point 34) had a significantly different number of OTUs from initial inoculum (time point 0).

**Figure 2.**
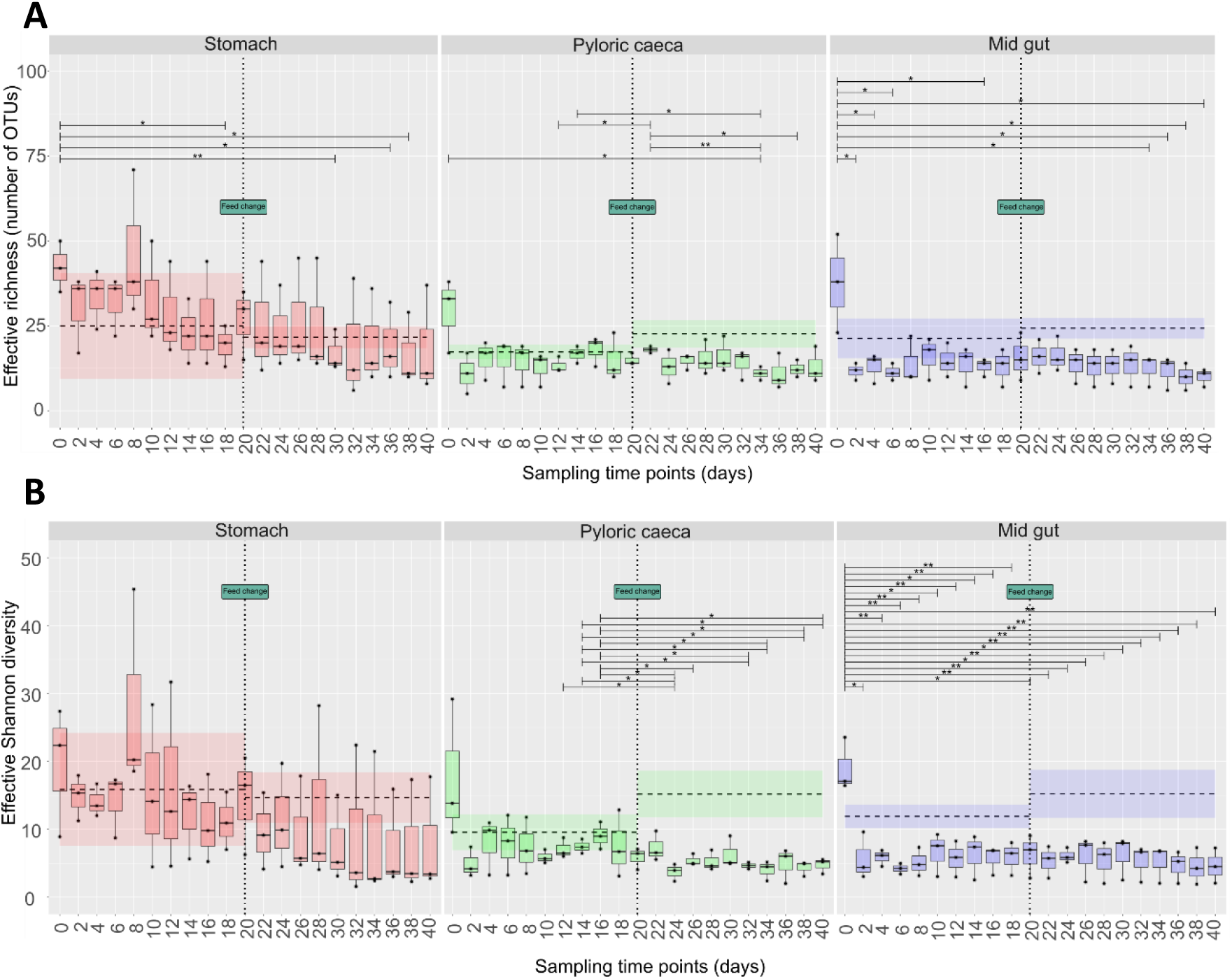
Calculated alpha-diversity metrics within SalmoSim system over time. The figure represents different alpha diversity outputs at different sampling time points (days) from SalmoSim system. Time point 0 represents microbial community composition within initial SalmoSim inoculum from the real salmon, time points 2-20 identifies samples from SalmoSim system fed on Fish meal diet, and time points 22-40 identifies samples from SalmoSim system fed on Fish meal free diet. The dashed line between days 0-20 represents average alpha diversity values measured in real salmon fed on Fish meal diet and dashed line between days 22-40 represents average alpha diversity values measured in real salmon fed on Fish meal free diet. Finally, the shaded region around the dashed line represents the standard deviation of the values measured within real salmon samples fed on the different diets. **A** visually represents effective richness (number of OTUs), **B** represents effective Shannon diversity and **C** represents effective Simpson diversity. The lines above bar plots represent statistically significant differences between different time points. The stars flag the levels of significance: one star (*) for p-values between 0.05 and 0.01, two stars (**) for p-values between 0.01 and 0.001, and three stars (***) forp-values below 0.001.

Figure 2 B indicated that within the stomach compartment effective Shannon diversity between all time points (including initial inoculum) was statistically similar (no statistical differences). However, within the midgut compartment, Shannon diversity metric was found to be statistically lower between time point 0 (initial inoculum) and the rest of the time points (sampling days 2-40). Finally, within pyloric caeca compartment it was found that effective Shannon diversity index was statistically different between time point 12 and 24, between time point 14 and time points 24, 32, 34, 38 and 40, between time point 16 and time points 24, 26, 32, 34, 38 and 40.

Taken together, diversity and richness estimates suggest some loss of microbial taxa as a result of transfer into SalmoSim in the pyloric caecum and midgut, but not in the stomach. Richness and diversity are then fairly stable over the time course of the experiment in stomach and mid gut compartments (some instabilities seen only between initial inoculum and later time points), whereas more instability in richness and diversity was seen in pyloric caeca compartment.

To assess the stability of the system inter-time point comparisons were undertaken examining with reference to pairwise beta-diversity metrics. Significant differences between time points represent instability in the system. Figure 3 visually summarises in-between time point comparison between different phylogenetic distances within SalmoSim system: unweighted UniFrac (Figure 3 A), balanced UniFrac (Figure 3 B), and weighted UniFrac (Figure 3 C). The comparison of all these different phylogenetic distances between different time points identified that only time point 2 was statistically different between the majority of the later time points. The general trend observed indicated that all gut compartments became increasingly stable throughout the 40-day experiment, with little-observed impact of introducing the different feed at day 20. This trend was supported by our qPCR data, suggesting increasing stability over the course of the 40-day experiment (Supplementary Figure 2).

**Figure 3.**
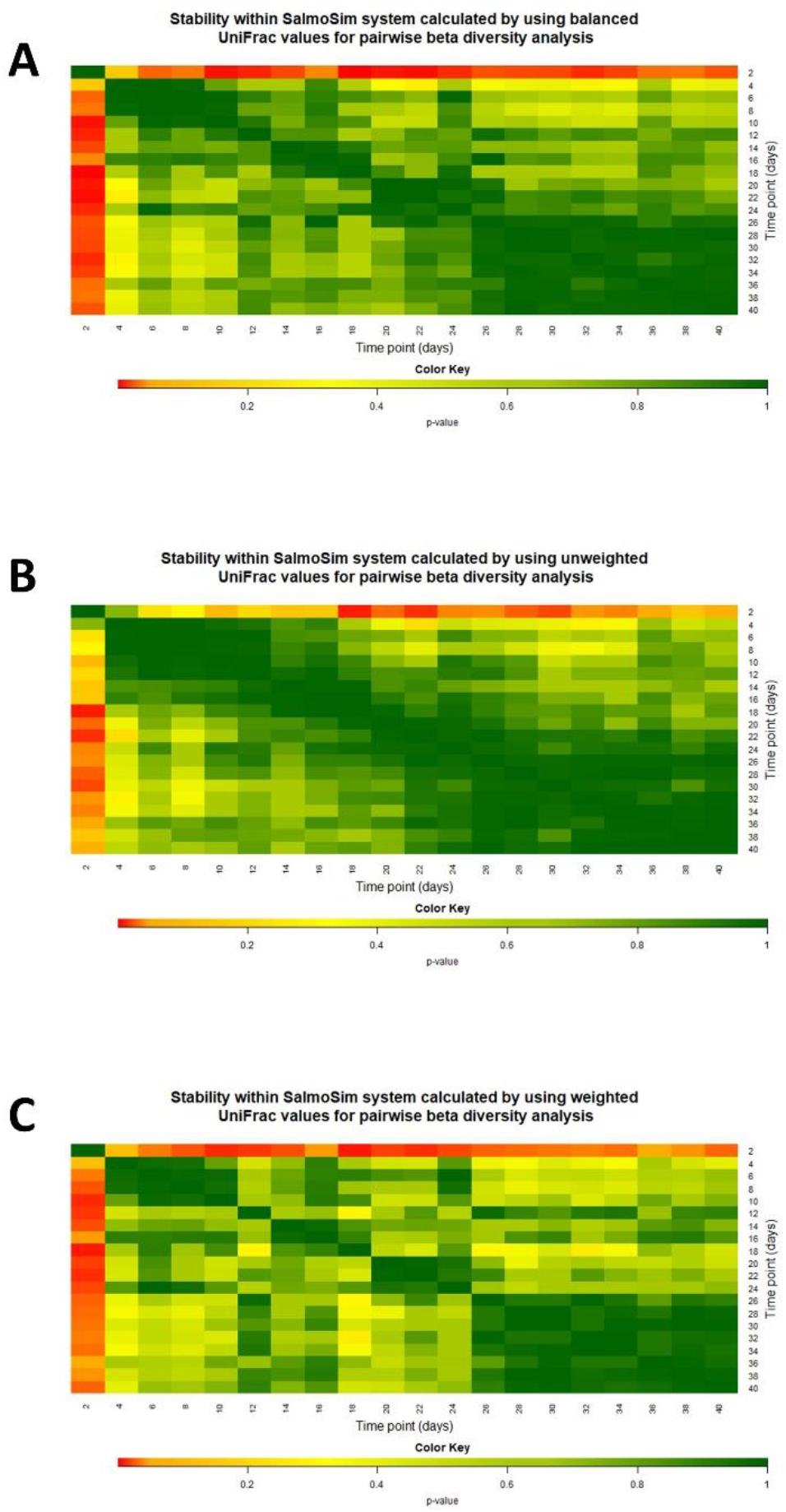
Stability within SalmoSim system calculated by using different UniFrac values for pairwise beta diversity analysis. The figure visually represents the microbial stability within the SalmoSim system (data from all gut compartments combined) as the pairwise beta diversity comparison between different sampling time points (days), calculated by using **A** unweighted (0%), **B** balanced (50%) and **C** weighted (100%) UniFrac as a distance measure. A small p-value indicates that the two time points are statistically different, and p>0.05 indicates that two time points are not statistically different. The colour key illustrates the p-value: red end of spectrum denoting low p values (low correlation between time points) and dark green indicating high p values (high correlation between timepoints).

### Comparisons of microbial identity and diversity between SalmoSim, starting inocula and real salmon

In order to compare different samples (inoculum, real salmon, SalmoSim) sample sizes were balanced by examining a reduced dataset that contained: real salmon samples fed on Fish meal diet, all inoculum samples and stable SalmoSim time points fed on Fish meal diet (days 36, 38, and 40). Alpha diversity comparisons are shown in Figure 2.

Several pairwise beta-diversity metrics were implemented to establish whether microbial compositions diverged between different sample types (real salmon, inoculum and SalmoSim-Table 2)). To explore the impact of rare OTUs in accounting for observed differences between sample types, several sub-setted datasets (listed in Table 2) were analysed using these beta-diversity metrics. Standard ecological metrics – Bray Curtis and Jaccard’s distance, that address the abundance and diversity of taxa in each community, did not identify significant differences between the inoculum and SalmoSim. In contrast, metrics that incorporate phylogenetic differences between taxa (i.e. Unifrac) did identify differences, indicating that there is variability between the inoculum and SalmoSim in terms of closely related taxa. Progressive removal of rare OTUs increased the compositional similarity of the inoculum and SalmoSim.

**Table 2.**
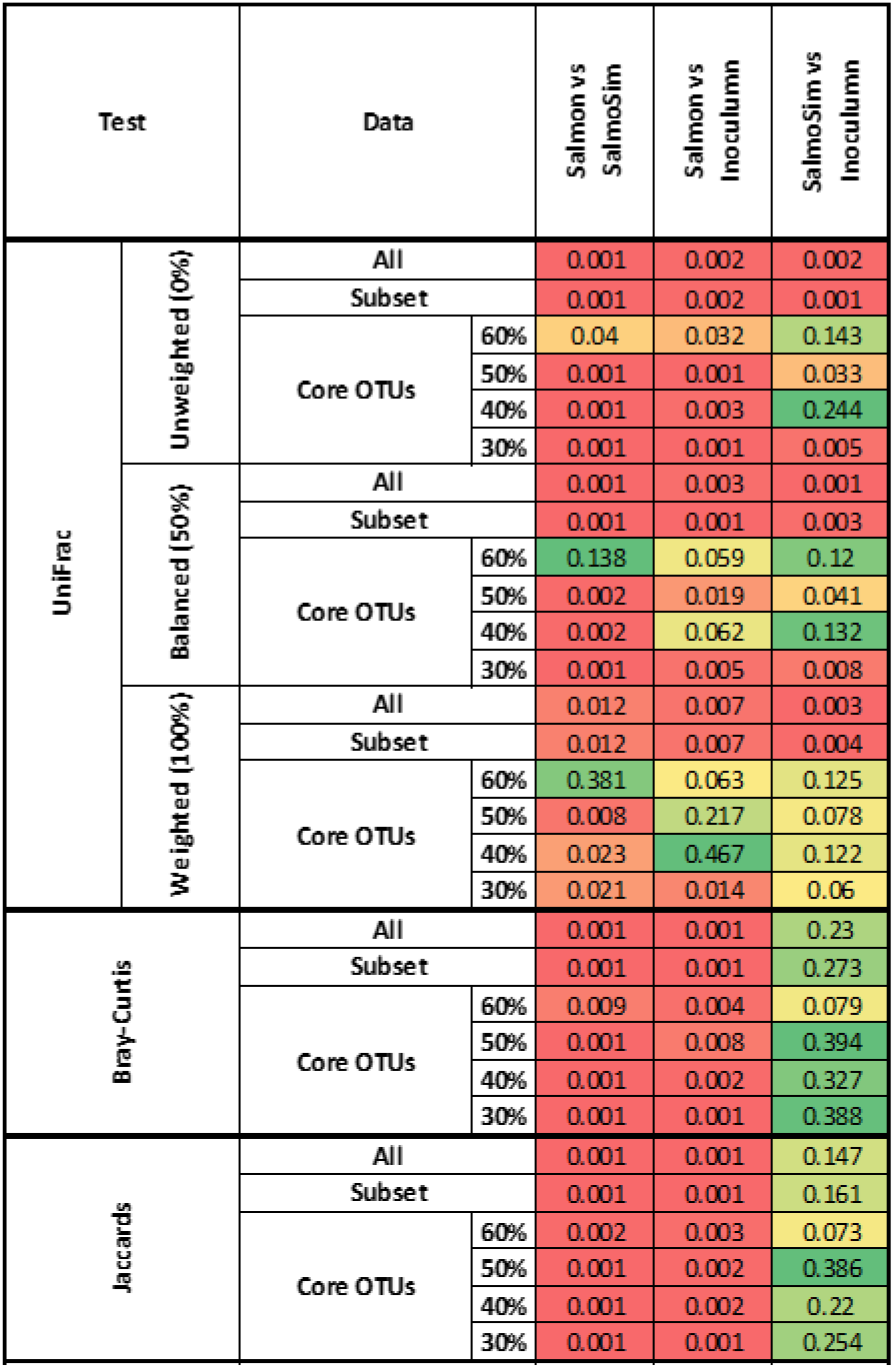
Beta diversity comparisons of microbial composition between different samples (real salmon, inoculum and SalmoSim) The table summarises different beta-diversity analysis outputs calculated by using different distances: phylogenetic (unweighted, balanced and weighted UniFrac) and ecological (Bray-Curtis and Jaccard ‘s), between different samples (data from all gut compartments combined): real salmon (**Salmon**), SalmoSim inoculum from the real salmon (**Inoculum**) and SalmoSim (only stable time points: 16, 18 and 20 fed on Fish meal diet, and 36, 38 and 40 fed on Fish meal free diet). A permutational multivariate analysis of variance (PERMANOVA) by using phylogenetic and ecological distances was performed to determine if the separation of selected groups is significant as a whole and in pairs. Numbers represent p-values, with p-values <0.05 identifying statistically significant differences between compared groups. The comparisons are shown for 3 different datasets: All (completed data set containing all the OTUs sequenced), Subset (containing OTUs that appear only in more than 3 samples and contribute to 99.9% of abundance within each sample), and core OTUs (containing OTUs that appear in 60%, 50%, 40% and 30% of the samples).

### Comparisons of microbial communities present within different gut compartments in SalmoSim and real salmon

Differences in the abundances of taxa measured using qPCR in each SalmoSim gut compartment were estimated running statistical methodologies described in Supplementary methods 1.2. Result shown in Supplementary Table 5 indicate that the pyloric caecum and midgut displayed the greatest similarity in community composition and that generally communities diverge more the further along the artificial alimentary canal they occur. Differences in rates of protein metabolism where observed between all gut compartments (Supplementary Table 5).

No divergence was observed in terms of alpha diversity between different gut compartments in both real salmon and SalmoSim samples (Supplementary Figure 3). In terms of microbial community composition, beta-diversity estimates of community differentiation also clearly identified that there was no statistical differences between different gut compartments in real salmon and SalmoSim based on both ecological (Bray-Curtis and Jaccard’s) and phylogenetic (UniFrac) distances (Supplementary Table 6).

### Effect of changing feed on the microbiome of real salmon by comparison to SalmoSim

#### The impact of feed on the abundance of individual taxon abundance

In order to investigate how bacterial groups within a different system (SalmoSim or real salmon) react to the change in feed, several statistical analyses were performed, initially on qPCR data (see Supplementary methods 1.3.). Table 3 summarises trends from this analysis and shows that in real salmon, the amount of Bacteroidetes decreased in all gut compartments in response to the change in the feed while the amount of all other bacterial groups did not change. By comparison, in the SalmoSim system several changes were observed in response to the feed change: Alphaproteobacteria decreased in stomach compartment, but increased in pyloric caeca and midgut compartments, Bacteroidetes decreased in stomach compartment, Firmicutes increased in all gut compartments, Gammaproteobacteria decreased in the stomach compartment, and Lactobacillus increased in stomach and midgut compartments, while the amount of all other bacterial groups remained not affected by the change in feed. 16S rRNA sequence-based comparisons of taxon abundance dynamics in salmon vs SalmoSim also indicate some differences as well as multiple similarities in the responses of the two systems to the feed trial (Figure 4 D). In this respect, the vast majority of OTUs (SalmoSim – 97%; Salmon – 95%; Figure 4 C) were unaffected by the change in feed; these included 161 OTUs shared by SalmoSim and the real salmon assayed. This is supported by Figure 5, which visually indicates that change in feed does not drastically affect the microbial composition in both real salmon and SalmoSim system. For OTUs whose individual abundance was impacted by feed across the two systems, only a single common OTU changed in the same way in both Salmon and SalmoSim (Figure 4 A). At higher taxonomic levels, 16S rRNA sequence data present a mixed picture (Figure 4 D), with broadly the same microbial taxa experiencing change in both salmon and SalmoSim (Bacteroidetes, Firmicutes, Gammaproteobacteria) but in no discernible overall pattern in either system.

**Figure 4.**
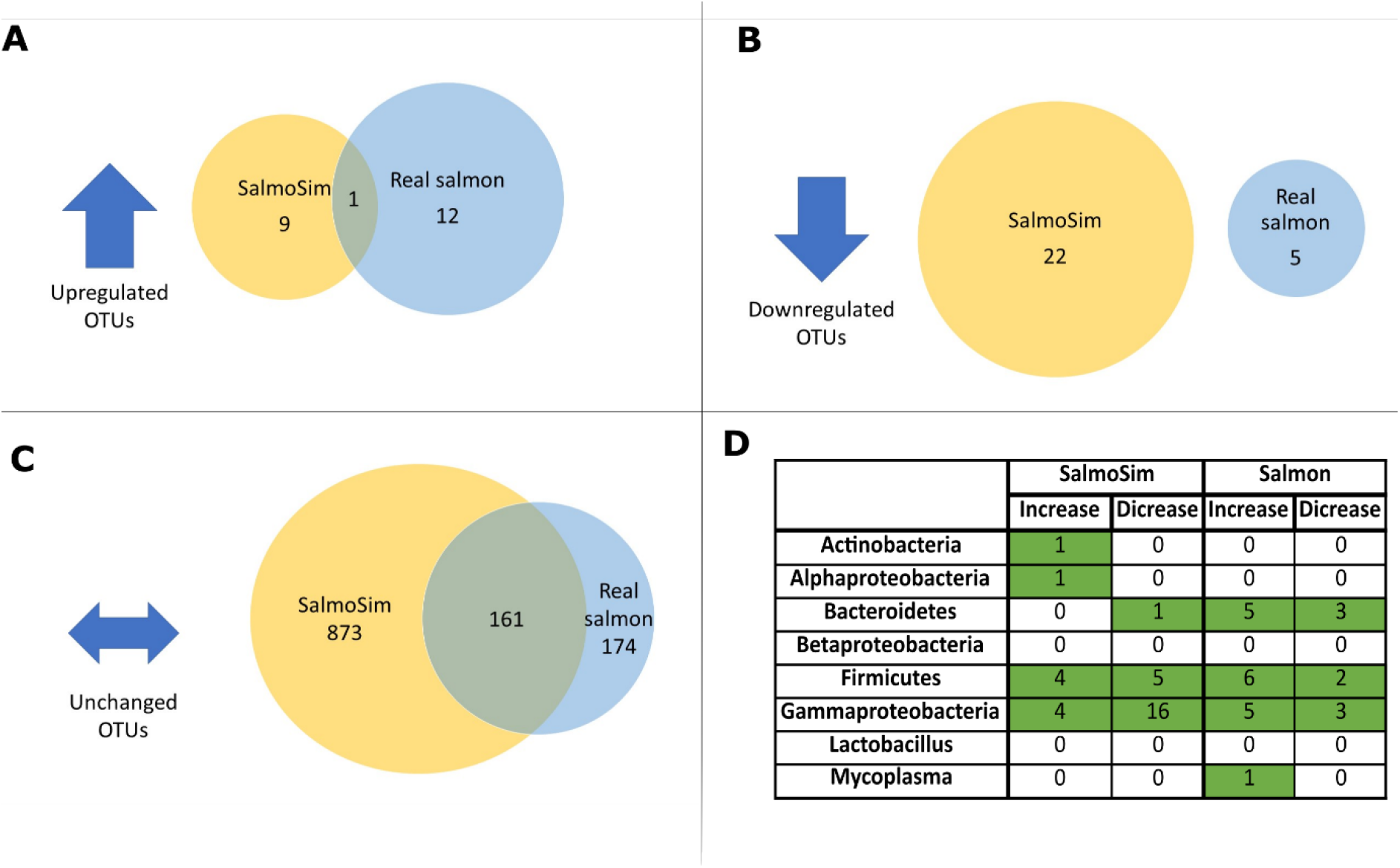
Differential abundance of OTUs within the real salmon and SalmoSim samples fed on Fish meal and Fish meal free diets. Figure visually represents differential abundance analysis output within the real salmon and SalmoSim samples fed on Fish meal and Fish meal free diets (data from all gut compartments combined). **A**: Venn diagram representing number of OTUs that were upregulated in both SalmoSim and real Salmon samples once the feed was switched, **B**: Venn diagram representing number of OTUs that were downregulated in both sample after the feed change, **C**: Venn diagram representing number of OTUs that did not change within SalmoSim and real salmon samples despite feed switch, **D**: table summarising number of OTUs that increased/decreased after feed change in real salmon and SalmoSim samples within different bacterial groups (that same that were analysed by using qPCR approach). Green colour indicates the values that are higher than 0.

**Table 3.** Summary of Estimated Marginal Means output for each mixed-effect linear model run with different measured values identifying the difference between real salmon and SalmoSim response to feed change (Fish meal to Fish meal free diet) The table identifies the trends in which measured qPCR values of bacterial groups were affected by changing feed (From fish meal to Fish meal free diet) between within two different samples (real Salmon and SalmoSim). Green cells identify no change in the bacterial group (p>0.05 from table 39) and blue values identify change (increase or decrease) in the bacterial group after the feed change from positive to negative (p<0.05 from table 40). **Bold** names identify similarities between SalmoSim and real salmon samples. The SalmoSim values used for this test involves stable SalmoSim time points: days 16, 18 and 20 (Fish meal diet), and days 36, 38 and 40 (Fish meal free diet).

|  | real salmon |  |  | SalmoSim |  |  |
| --- | --- | --- | --- | --- | --- | --- |
|  | S | PC | MG | S | PC | MG |
| **Actinobacteria** | no change | no change | no change | no change | no change | no change |
| **Alphaproteobacteria** | no change | no change | no change | decrease | increase | increase |
| **Bacteroidetes** | decrease | decrease | decrease | decrease | no change | no change |
| **Betaproteobacteria** | no change | no change | no change | no change | no change | no change |
| **Firmicutes** | no change | no change | no change | increase | increase | increase |
| **Gammaproteobacteria** | no change | no change | no change | decrease | no change | no change |
| **Lactobacillus** | no change | no change | no change | increase | no change | increase |
| **Mycoplasma** | no change | no change | no change | no change | no change | no change |

#### Sequence-based assessments of microbial alpha and beta-diversity diversity in SalmoSim and real salmon

Figure 5 visually represents the microbial composition within different gut compartments in real salmon and SalmoSim system. Figure 5 indicates that most gut compartments for both real salmon and SalmoSim are dominated by *Pseudomonas, Psychrobacter* and *Staphylococcus* genera and also suggest that genera present in the inoculum are generally maintained in SalmoSim. In terms of change in alpha diversity, the only statistically significant difference in response to the switch in feed was observed in the pyloric caeca compartment of the SalmoSim compartment based on the Shannon diversity metric (Supplementary Figure 4), where a slight decrease alongside the fishmeal free fish diet occurred. Otherwise, the change in feed formulation did not impact richness or diversity in any gut compartment, either in real salmon, or in SalmoSim.

**Figure 5.**
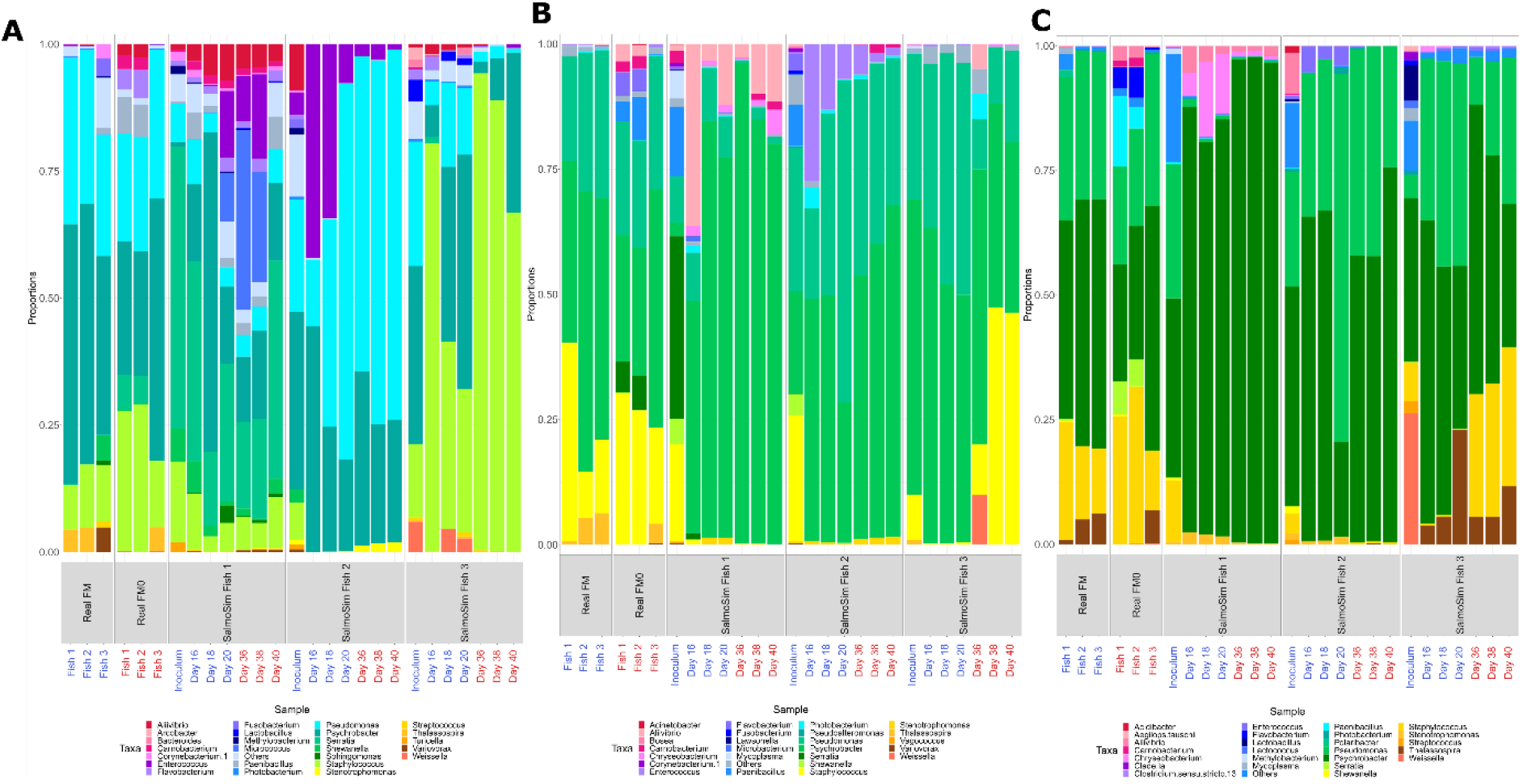
Microbial composition (25 most common genus + others) amongst sample types and feeds. **A**: microbial composition within stomach compartment, **B**: microbial composition within pyloric caeca compartment, and **C**: microbial composition within midgut compartment. The different sample types are represented by the labels on the x-axis: Real FM (real salmon fed on Fish meal), Real FM0 (real salmon fed on Fish meal free diet), SalmoSim Fish 1-3 (SalmoSim biological replicate runs 1-3). Labels in blue represent samples fed on Fish meal diet and in red samples fed on Fish meal free diet. For SalmoSim only stable time points for each feed were selected: time points 16-20 for Fish meal diet, and time points 36-40for Fish meal free diet.

Phylogenetic and ecological distance beta-diversity metrics were deployed and indicate that changing feed was a key driver of community composition in both salmon and SalmoSim (Figure 6 E). The effect was diminished when a lower number of putatively ‘stable’ SalmoSim time points were examined (Figure 6 E).

**Figure 6.**
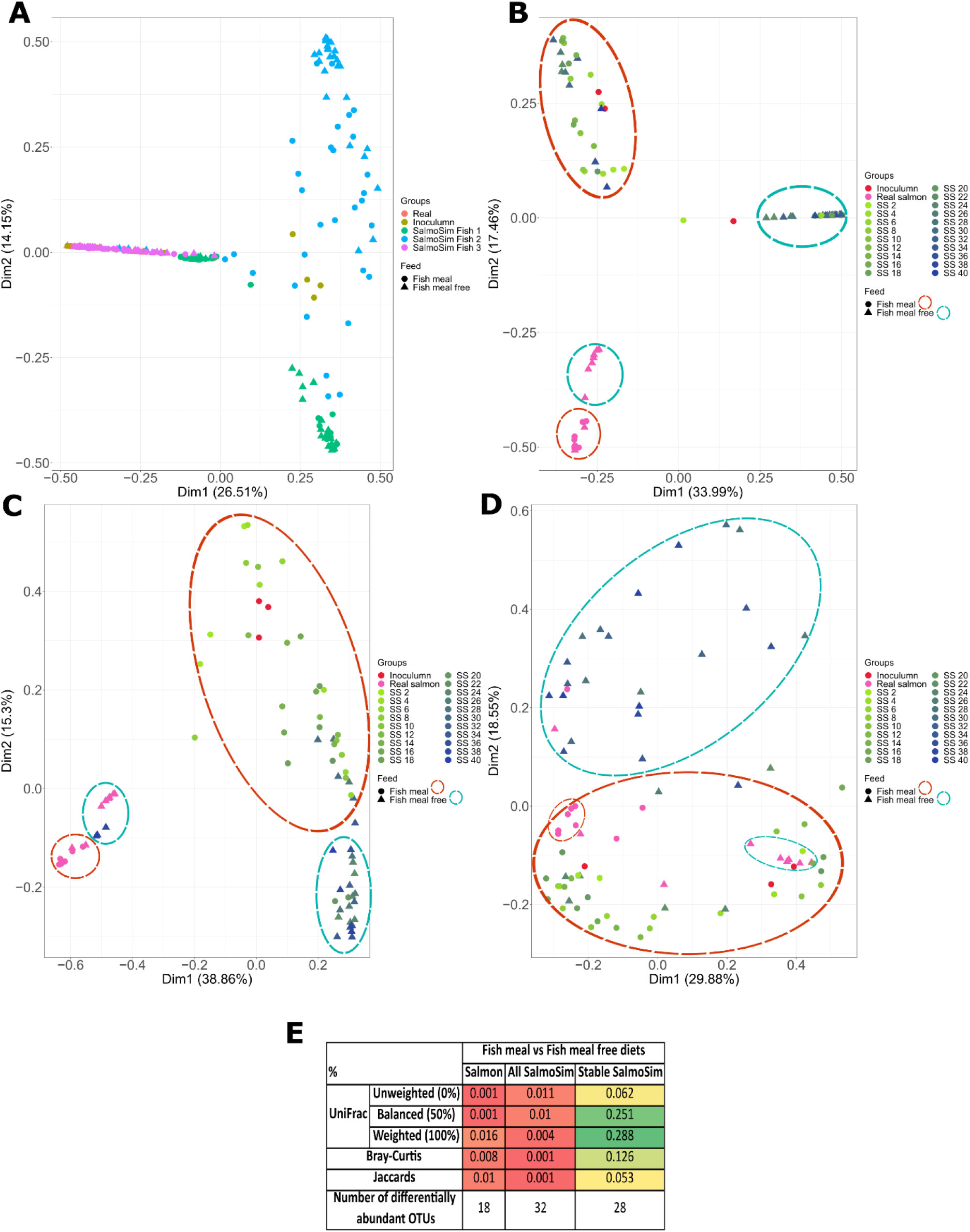
PCoA plots, beta diversity and differential abundance values visualising bacterial communities within different samples (SalmoSim and real salmon) and feeds (Fish meal and Fish meal free diets) Figure visualises four principal-coordinate analysis (PCoA) plots for Bray-Curtis dissimilarity measures for different samples (Inoculum, real salmon and SalmoSim), different sampling time points from SalmoSim system, different biological replicates and different feeds. **A** represents all sequenced data together in which different colours represent different samples (real salmon, inoculum and 3 different SalmoSim biological replicates (Fish 1, Fish 2, Fish 3)) and different shapes represent different feeds; **B-D** represent sequenced data together for real salmon, inoculum and different biological replicates of SalmoSim (**B**: Fish 1, **C**: Fish 2, **D**: Fish 3). In figures B-D different colours represent different samples (inoculum, real salmon and different sampling points of SalmoSim), different shapes represent samples fed on two different feeds, and samples fed on same feeds were circled manually in dotted circles. Dim 1 is principal coordinate 1, and Dim 2 is principle coordinate 2. Finally, table **E** summarises different beta-diversity analysis outputs calculated by using different distances: phylogenetic (unweighted, balanced, and weighted UniFrac) and ecological (Bray-Curtis and Jaccard’s), between samples fed on Fish meal or Fish meal free diets. Numbers represent p-values, with p-values <0.05 identifying statistically significant differences between compared groups. It also indicates the differential abundant number of OTUs between samples fed on Fish meal and Fish meal free diets. It also indicates the differential abundant number of OTUs between samples fed on Fish meal and Fish meal free diets. The comparisons are shown for four different datasets: Salmon (containing sequenced samples from real salmon), All SalmoSim (containing all samples from SalmoSim system), and Stable SalmoSim (containing samples only from stable time points: 16, 18 and 20 fed on Fish meal, and 36, 38 and 40 fed on Fish meal free diet).

To provide an overview of microbial composition and variation in the experiment, a PCoA plot was constructed based on Bray-Curtis distanced between samples (Figure 6 A-D). As with Figure 5, biological replicate (the founding inoculum of each SalmoSim run) appears to be a major driver of community composition in the experiment (Figure 6 A, Figure 5). Only once individual SalmoSim replicates are visualised separately do the changes to microbial communities in response to the feed become apparent (Figures 6 B-D) and reflect PERMANOVAs carried out in Figure 6 E. Inocula for the respective replicates cluster among SalmoSim samples for the fish meal diet in each case. Samples from real salmon fed on the different diets diverge from one and other (supported by Figure 6 E, Figure 5), however, not necessarily along the same axes as each SalmoSim sample indicative, perhaps, or a different microbiological basis for that change.

### Volatile Fatty Acid (VFA) production in SalmoSim during the feed trial

VFA levels were measured throughout the SalmoSim trial for the stable time points (time points 18 and 20 for Fish meal, and time points 38 and 40 for Fish meal free diet). These results are represented in Figure 7 that indicate that no significant differences in any VFA production by the system was noted in any of the gut compartments after the introduction of the plant-based feed. We did not also note any significant differences between different gut compartments apart from the amount of formic acid between stomach and mid gut compartments fed on Fish meal diet and the butanoic acid concentration between stomach and mid gut compartments fed on Fish meal free diet (Figure 7).

**Figure 7.**
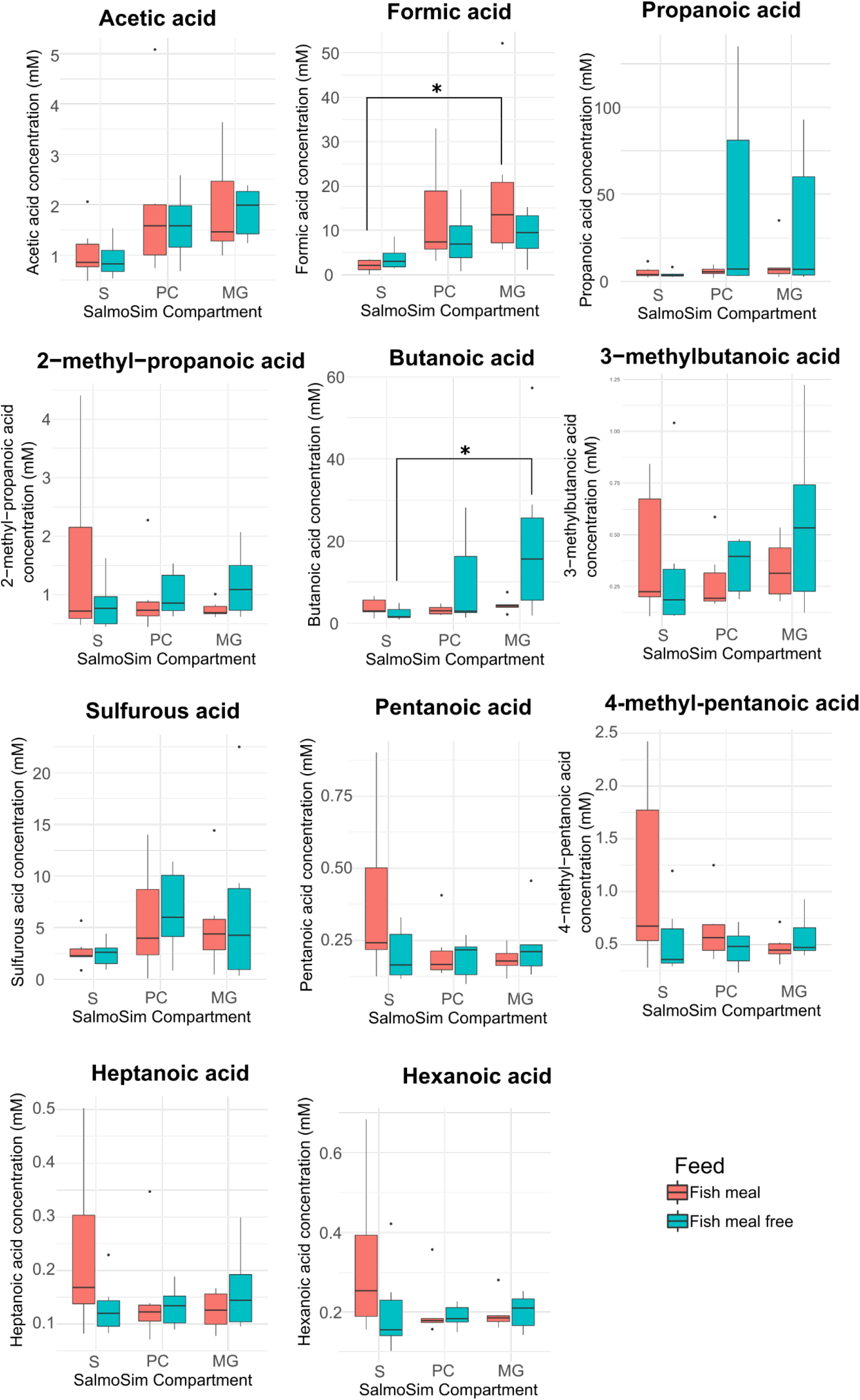
Volatile fatty acid production in the SalmoSim system fed on Fish meal and Fish meal free diets within different gut compartment. Figure visually represents 11 volatile fatty acid production within SalmoSim system fed on Fish meal and Fish meal free diets within different gut compartments. X axis represents the concentration of specific volatile fatty acid (mM) while the Y axis represents each gut compartment (stomach, pyloric caeca, midgut). Red colour denoted Fish meal and blue – Fish meal free diets. The lines above bar plots represent statistically significant differences between different feeds and gut compartments. The stars flag the levels of significance: one star (*) for p-values between 0.05 and 0.01, two stars (**) for p-values between 0.01 and 0.001, and three stars (***) for p-values below 0.001.

### Microbial correlates with SalmoSim fermentative profiles

Several potential physical correlates with microbial activity were measured from the SalmoSim system including a range of volatile fatty acids, total protein content of each compartment and the level of ammoniacal nitrogen present. To assay the potential microbial drivers of these profiles Pearson correlation coefficients across different values measured and OTUs. Table 4 indicates that the strong statistically significant negative and positive correlations were found between acetic acid and 8 (3 OTUs belonging to *Psychrobacter* genus, 3 to *Pseudomonas*, 1 to *Enterococcus*, and 1 OTU to *Aliivibrio*) and 1 (belong to *Pseudomonas* genus) OTUs respectively. This figure also shows that formic acid negative correlated to OTU belonging to Alviibrio genus, propanoic acid negative correlated with 2 OTUs belonging to *Psychrobacter* and *Pseudomonas* genus, and 3-methyl butanoic acid negatively correlated to OTU belonging to *Psychrobacter* genus. Finally, 2-methyl propanoic acid positively correlated to OTU belonging to *Enterococcus* genus.

**Table 4.**
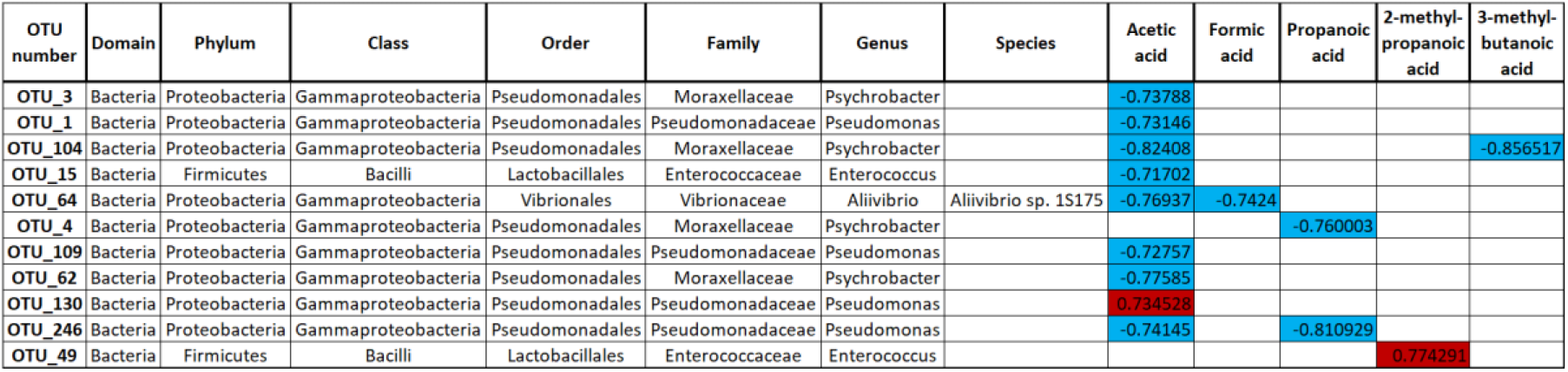
Person correlation coefficients across different values measured and taxonomic variables. Table summarises calculated statistically significant (p<0.05) and strongly correlated (r>0.7) Pearson correlation coefficients across a set of meta- and taxonomic variables. Blue colour represents positive correlation and red colour represents negative correlations. Numbers in cells represent r values with 1 being strong positive correlation and −1 – strong negative correlation.

## Discussion

Our findings suggest some loss of microbial taxa diversity and richness as a result of transferring initial inoculums from real salmon into the SalmoSim system in the pyloric caeca and mid gut compartments. Several lines of evidence suggest that rare taxa make up the majority of those lost, and progressive removal of rare OTUs increased the compositional similarity of the inoculum and SalmoSim via all metrics. Nonetheless, comparisons of microbial diversity suggested no significance differences between the composition of initial inoculum and SalmoSim system based on standard ecological metrics without the removal of any OTUs. A general trend was observed in which all gut compartments became increasingly stable throughout the 40-day experiment, with little-observed impact of introducing the different feed at day 20. Unlike for the inoculums, comparisons of real salmon and SalmoSim samples at the microbial level showed significant differences using both ecological and phylogenetic metrics. This could be explained by the fact that samples used for real salmon and SalmoSim originated from different individuals, whereas initial inoculum and SalmoSim samples for a given run originated from the same fish. While we saw, correspondingly, that the biological replicate (the founding inoculum of each SalmoSim run) was the major driver of community composition in the experiment, once the individual runs were separated, the phylogenetic and ecological beta-diversity metrics suggested that changing feed was a driver of community composition in both real salmon and SalmoSim. However, the vast majority of OTUs remained unchanged by the switch in feed in both real salmon and SalmoSim and no changes were indicated in the bacterial activity (VFA production) within the system after the introduction of plant-based feed.

Many of the microbes we detected, and cultured, from the salmon gut microbiome have been reported previously from this species. For example, gram-negative *Pseudomonas* and *Psychrobacter*, the most abundant genera we observed, are among the core bacterial taxa known to reside within the real salmon gut (Gajardo et al., 2016; Navarrete et al., 2009; Webster et al., 2018). *Staphylococcus* genera have also been reported widely in fresh-water and marine salmon (Dehler et al., 2017). SalmoSim was able to maintain these species in culture throughout the experimental run, and although some diversity was lost, no statistical differences could be detected between the composition of SalmoSim and the fish gut communities used to found the different biological replicates. Notable by their scarcity were mycoplasma OTUs, which occurred at relatively low abundance in both the *in vivo* and *in vitro* systems in this study. *Mycoplasma* OTUs were recovered from most SalmoSim gut compartments at low abundances (see supplementary Table 7), suggesting that these fastidious microbes can survive in the bioreactors. Our group and several others have widely reported *Mycoplasma* species from marine and freshwater stage Atlantic salmon, where many proliferate intracellularly in the gut epithelial lining (Cheiab *et al*., unpublished; (Heys et al., 2020; Llewellyn et al., 2014). Establishing whether mycoplasma can actively proliferate in SalmoSim would require the use of founding communities rich in these organisms in a future experiment, and we found, as many others working with microbiomes do (Jones et al., 2018), that interindividual variability (in our case affecting the initial inoculum) was a main driver for gut microbial composition divergence.

Previous attempts to map compositional differences between the microbial communities of salmon gut compartments indicate significant divergences (Gajardo et al., 2017b; Heys et al., 2020). We failed to detect significant differences between stomach, pyloric caecum and midgut of the fish sampled in this experiment, and the same is true for SalmoSim when the *in vivo* / *in vitro* data are matched for sample size (N=3). Increasing the number of SalmoSim timepoints included in this comparison does result in significant divergence emerging *in vitro*, consistent with previous *in vivo* studies, and perhaps reflecting our relatively small *in vivo* sample size.

We identified that a change in feed resulted in an overall shift in microbial community structure in both real salmon and SalmoSim system, as was also found to be the case in many previous studies (Egerton et al., 2020; Gajardo et al., 2017a; Michl et al., 2017). The direction of this shift, and the microbial taxa involved, were not equivalent in SalmoSim and real salmon, although no overall trend was observed at higher taxonomic levels in either system Importantly, it is also the case that the vast majority of OTUs within both real salmon and SalmoSim were not affected by the switch in feed. Furthermore, it was found that change in feed did not affect VFA production in the SalmoSim system. As such, it is not clear whether any relevant functional shifts occurred in the microbiome of SalmoSim or real salmon as a result of the treatment. This lack of change is not unexpected, considering the plant-based feed was developed to have similar nutritional composition to a Fish meal-based feed.

The use of *in vitro* systems to study and model the microbial communities of monogastric vertebrates is becoming increasingly widespread, simulating: pig (Tanner et al., 2014), chicken (Card et al., 2017), dog (Duysburgh et al., 2020) and other vertebrate guts. For example, using *in vitro* gut simulators is a widely accepted approach to study the human gut microbiome (Déat et al., 2009; Kim et al., 2016; Van Den Abbeele et al., 2010). One of the most established systems is the Simulator ofthe Human Intestinal Microbial Ecosystem (SHIIME) that mimics the entire gastrointestinal tract incorporating stomach, small intestine and different colon regions (Molly et al., 1994). This system was used to study the effects of many different dietary additives on human microbiome (Giuliani et al., 2016; Sánchez-Patán et al., 2015). The value of *in vitro* simulators in providing genuine insights is limited by the research question and the corresponding level of sophistication required. The host component of the system, for example, is often poorly modelled, although cell lines, artificial mucosae and digestion / absorbance systems can be included, to provide specific insights (Déat et al., 2009; Van den Abbeele et al., 2012). As we found, inter-individual variability may be an important consideration, and adequate biological replication is necessary to enable reliable interpretation of results, a consideration that can be overlooked even by the most sophisticated systems. Prior to the current study, only one other attempt was made to study the effect of diet on Atlantic salmon gut microbial composition *in vitro* (Zarkasi et al., 2017). In this preliminary study a simple *in vitro* system was used to assess the impact of different feed formulations on the microbial communities of faecal slurries prepared from live salmon. However, no direct comparison was made with a true *in vivo* trial; nor were the different gut compartments present in salmon modelled in any detail and the predictive value for such simple *in vitro* systems in not immediately clear. Nonetheless, the work provided an important catalyst for the development of most sophisticated systems.

Our results indicate that SalmoSim could not only maintain stable microbial communities from real salmon, but also demonstrates similar responses to experimental treatment as those seen in real salmon. These results are encouraging, however, the nature of the treatment applied in this study: a switch between two similar feeds that had little effect on the gut microbiota *in vivo*; suggests that further experimentation with SalmoSim would be beneficial. For example, the survival and influence of probiotics within the system or the influence of known prebiotics could also be assessed, as they have in other *in vitro* gut systems (Duysburgh et al., 2020). Gut models such as SalmoSim could have a powerful role in aquaculture, where there is intense innovation around feed and feed additives (Encarnação, 2016; Hartviksen et al., 2014; Kristiansen et al., 2011), while capacity for *in vivo* trials is limited. The aim of such systems could be to provide pre-screening tool for new feed ingredients and additives with the aim of reducing the cost and scale of *in vivo* testing. In parallel, an *in vitro* gut model for salmon could also be exploited to understand questions of public health importance (e.g. antimicrobial resistance and transfer (Card et al., 2017), or the fundamental ecological processes that underpin microbiome dynamics and assembly.

