## Supplemental Methods for "Development of a three-compartment *in vitro* simulator of the Atlantic Salmon GI tract and associated microbial communities: SalmoSim"

Supplementary methods

### 1. qPCR data analysis

#### 1.1. Investigating bacterial dynamics within SalmoSim system over time

The subdivided datasets by different SalmoSim gut compartment and different feed were used to run Model 1 with different values measured during a validation experiment considering run (biological replicates) as a random effect. The normality and heterogeneity of the residuals for of each model were checked by using Shapiro and Bartlett's tests. If these tests showed that residues are not normally distributed or not heterogeneous, the data was transformed by using Code 1 and then used to re-run Model 1. Finally, the post-hoc test Estimated Marginal Means (also known as Least-Squares Means) was used Model 1 ran with different values in order to investigate in-between individual time point comparison.

Model 1 = lme(value~ Time, random =c(~1|Run), data=**groupX**)

**Model 1 Mixed effect linear model formula to investigate the effect of time taking run as a random effect.** *In Model 1 groupX denotes subdivided data sets by different SalmoSim compartments (stomach, pyloric caecum or midgut), and value denotes the qPCR results for one of the different targeted bacterial groups (Actinobacteria, Alphaproteobacteria, Bacteroidetes, Betaproteobacteria, Firmicutes, Gammaproteobacteria, Lactobacillus, Mycoplasma) or protein or ammonia concentrations. This model takes different SalmoSim runs as a random effect (biological replicates).*

```
if(lambda!=0){y=((value)^lambda-1)/lambda}

if(lambda==0){y=log(value)}
```

**Code 1 Code used to transform the data.** *Value in this code denotes the qPCR results for one of the different targeted bacterial groups (Actinobacteria, Alphaproteobacteria, Bacteroidetes, Betaproteobacteria, Firmicutes, Gammaproteobacteria, Lactobacillus, Mycoplasma) or protein or ammonia concentrations. Lambda values in 0.1 increments were tested.*

#### 1.2. Comparison between different SalmoSim compartments

The full datasets containing all three biological replicates was used to run Model 2 that considers run (biological replicates) and time as random effects. The normality and heterogeneity of each model residues were checked by using Shapiro and Bartlett's tests. If these tests showed that residues are not normally distributed or not heterogeneous, the data was transformed by using Code 1 and then used re-run Model 2. Finally, post-hoc test Estimated Marginal Means (also known as Least-Squares Means) test was applied to each model run with different transformed or not transformed values to investigate statistical similarities/differences between different SalmoSim reactor vessels.

Model 2 = lme(value~ Compartment, random =c(~1|Run,~ 1|Time))

**Model 1 Mixed effect linear model formula to investigate the effect of SalmoSim compartment taking run (biological replicates) and time as a random effect.** *In Model 2, value denotes the qPCR results for one of the different targeted bacterial groups (Actinobacteria, Alphaproteobacteria, Bacteroidetes, Betaproteobacteria, Firmicutes, Gammaproteobacteria, Lactobacillus, Mycoplasma) or protein or ammonia concentrations. This model takes run and time as random effects.*

#### 1.3. Comparing *in vivo* and *in vitro* trials

The subdivided datasets for each gut compartment for both real salmon (all samples) and SalmoSim (samples from only stable time points) were input into Model 3. In order to investigate how bacterial groups within different type of sample (SalmoSim or real salmon) react to the change in feed Post-hoc test Estimated Marginal Means (also known as Least-Squares Means) was used in order to have a more detailed look at the effect of the interaction between feed and sample on the abundance of each target taxon.

Model 3 = lm(value~ Feed*samples, data=GroupX)

**Model 3 Linear model formula to investigate the effect of interaction between feed (Fish meal and Fish meal free diets) and sample (real salmon and SalmoSim samples) on the qPCR values measured.** *In model 3, value denotes the qPCR results for one of the different targeted bacterial groups (Actinobacteria, Alphaproteobacteria, Bacteroidetes, Betaproteobacteria, Firmicutes, Gammaproteobacteria, Lactobacillus, Mycoplasma). Group X is a subset of dataset separated by different gut compartments (stomach, pyloric caecum and midgut). Feed identifies Fish meal and Fish meal 0 diets, and sample identifies real salmon and stable time point SalmoSim samples (days 16, 18 and 20 SalmoSim fed on Fish meal diet, and days 36, 38, and 40 SalmoSim fed on Fish meal free diet).*

#### 2. Volatile Fatty Acid (VFA) analysis

The measured VFA values were input to Model 4 that took time point (sampling time point) and run (biological replicate of SalmoSim system) as random effects. This was followed by the Post-hoc test Estimated Marginal Means (also known as Least-Squares Means) in order to have a more detailed look at the effect of the interaction between feed and SalmoSim compartment affect the concentration of VFAs.

**Model 4 = lmer(VFA~ Feed*Compartment+(1|Time point)+(1|Run))**

**Model 4 Mixed effect linear model to investigate the significance of different VFA concentrations between SalmoSim fed on Fish meal and Fish meal free diets within different SalmoSim compartments.** *In Model 4 VFA denotes the VFA values measured. This model takes run and time as random effects.*
