## Supplemental Figure 1 for "Development of a three-compartment *in vitro* simulator of the Atlantic Salmon GI tract and associated microbial communities: SalmoSim"

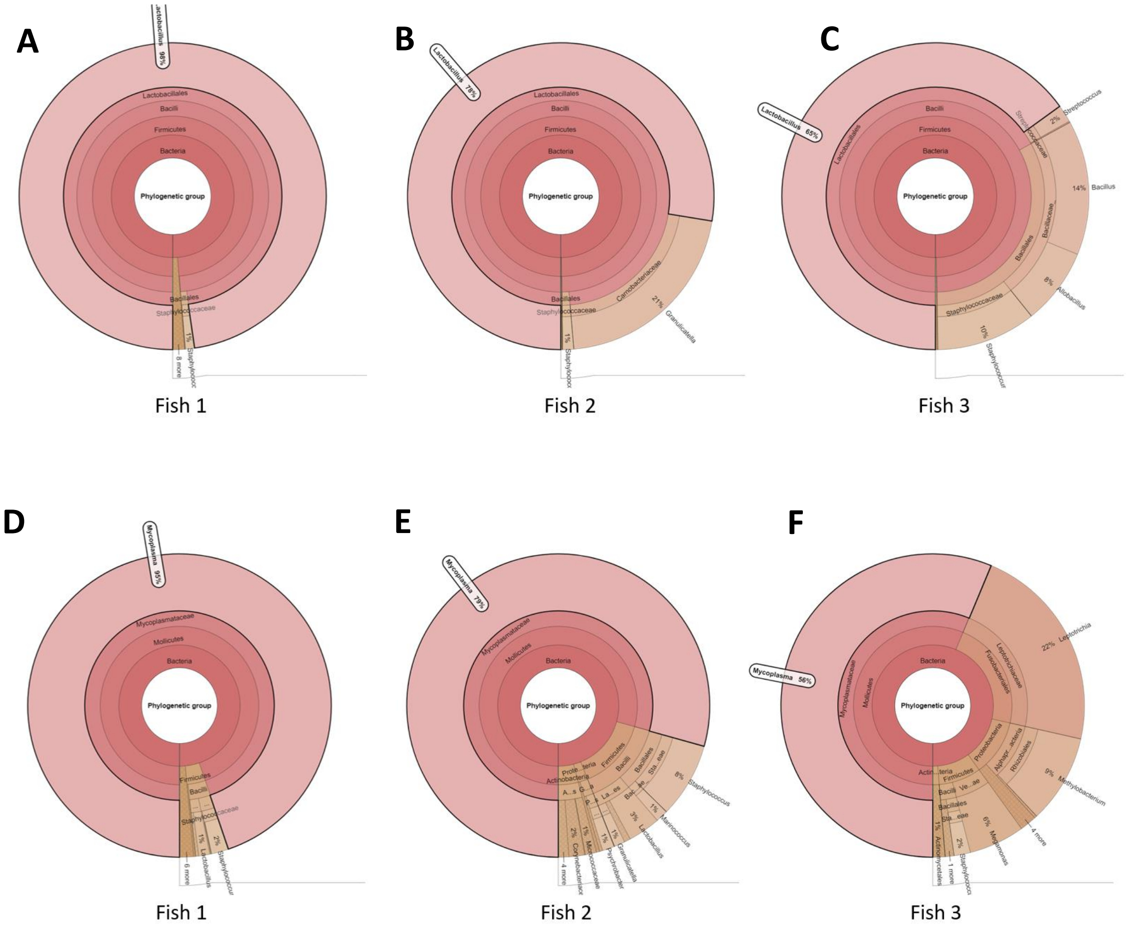


**<u>Figure 1</u> Specificity of the primers that target Lactobacillus and Mycoplasma genus** *The results in figure summarise bacterial genus targeted by Lactobacillus (Figure 1 A, B, C) and Mycoplasma (Figure 1 D, E, F) specific primer set. It shows that of all genus captured by Lactobacillus primer pair 98% were Lactobacillus in fish 1, 78% in fish 2, and 65% in fish 3. While of all genus captured by Mycoplasma primer pair 95% were Mycoplasma in fish 1, 79% in fish 2, and 56% in fish 3.*
