## Supplemental Figure 2 for "Development of a three-compartment *in vitro* simulator of the Atlantic Salmon GI tract and associated microbial communities: SalmoSim"

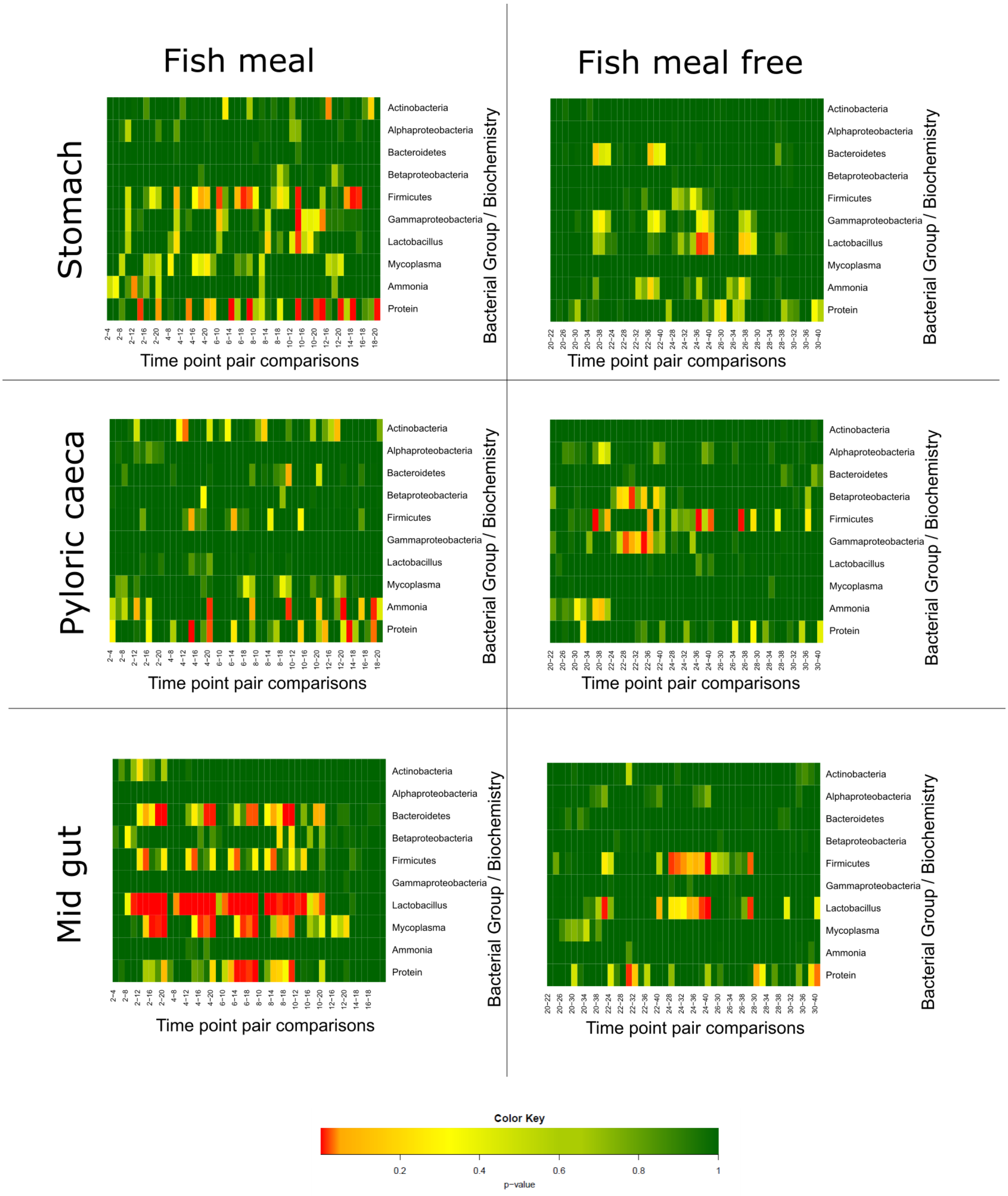


<u>**Figure 2**</u> **Measured value (qPCR, ammonia and protein concentrations) stability within different SalmoSim compartments fed on Fish meal and Fish meal free diets** *The figure summarises the Estimated Marginal Means output for each mixed-effect linear model (Model 1**Error! Reference source not found.**) run with different values measured in different SalmoSim compartments (qPCR measurements, ammonia and protein concentrations) identifying the difference between different time points during the first (system fed on Fish meal diet) and last 20 days (system fed on Fish meal free diet) of validation experiment. A small p-value indicates that the two time points are statistically different, and p>0.05 indicates that two time points are not statistically different. The colour key illustrates the p-value: red end of spectrum denoting low p values (low correlation between time points) and dark green indicating high p values (no differences between timepoints).*
