## Supplemental Figure 3 for "Development of a three-compartment *in vitro* simulator of the Atlantic Salmon GI tract and associated microbial communities: SalmoSim"

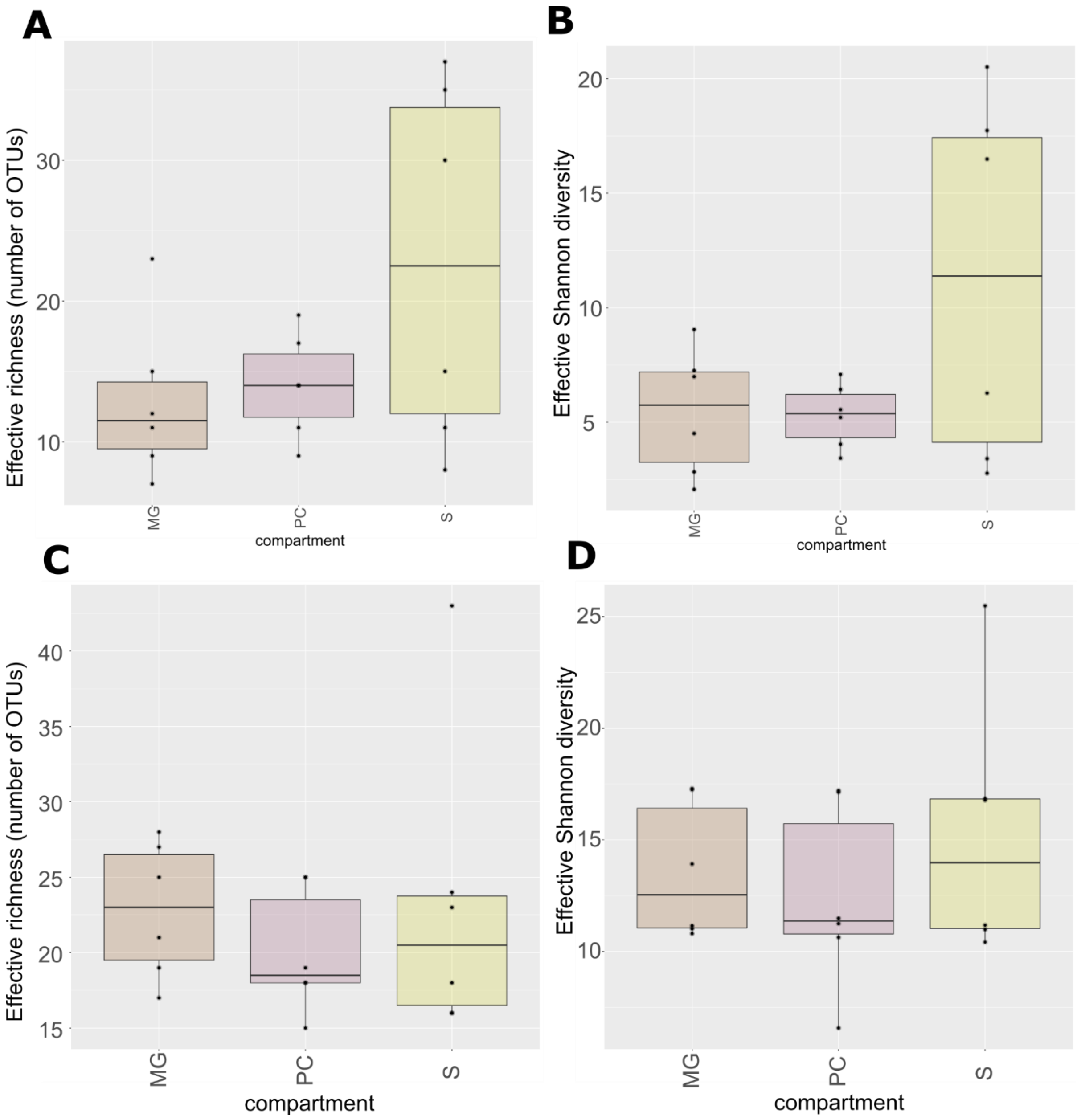


**<u>Figure 3</u> Calculated alpha-diversity metrics within different SalmoSim and real salmon gut compartments** *Figure shows different alpha diversity outputs in different SalmoSim (**A** and **B**) and real salmon (**C** and **D**) gut compartments. Equal number of samples (n=18) for each dataset were selected: real salmon (samples from the 3 gut compartments, from 3 biological replicates, for each of the 2 feeds) and SalmoSim (samples from each of the 3 gut compartments, for 3 biological replicate runs, for each of the 2 feeds (time point 20 for Fish meal and time point 40 for Fish meal free diet)). **A** and **C** visually represents effective richness (number of OTUs) and **B** and **D** represents effective Shannon diversity. **MG** denotes midgut compartment (red), **PC** – pyloric caeca (purple), and **S** – stomach (yellow). The lines above bar plots represent statistically significant differences between groups (gut compartments). The stars flag the levels of significance: one star (*) for p-values between 0.05 and 0.01, two stars (**) for p-values between 0.01 and 0.001, and three stars (***) for p-values below 0.001.*
