## Supplemental Figure 4 for "Development of a three-compartment *in vitro* simulator of the Atlantic Salmon GI tract and associated microbial communities: SalmoSim"

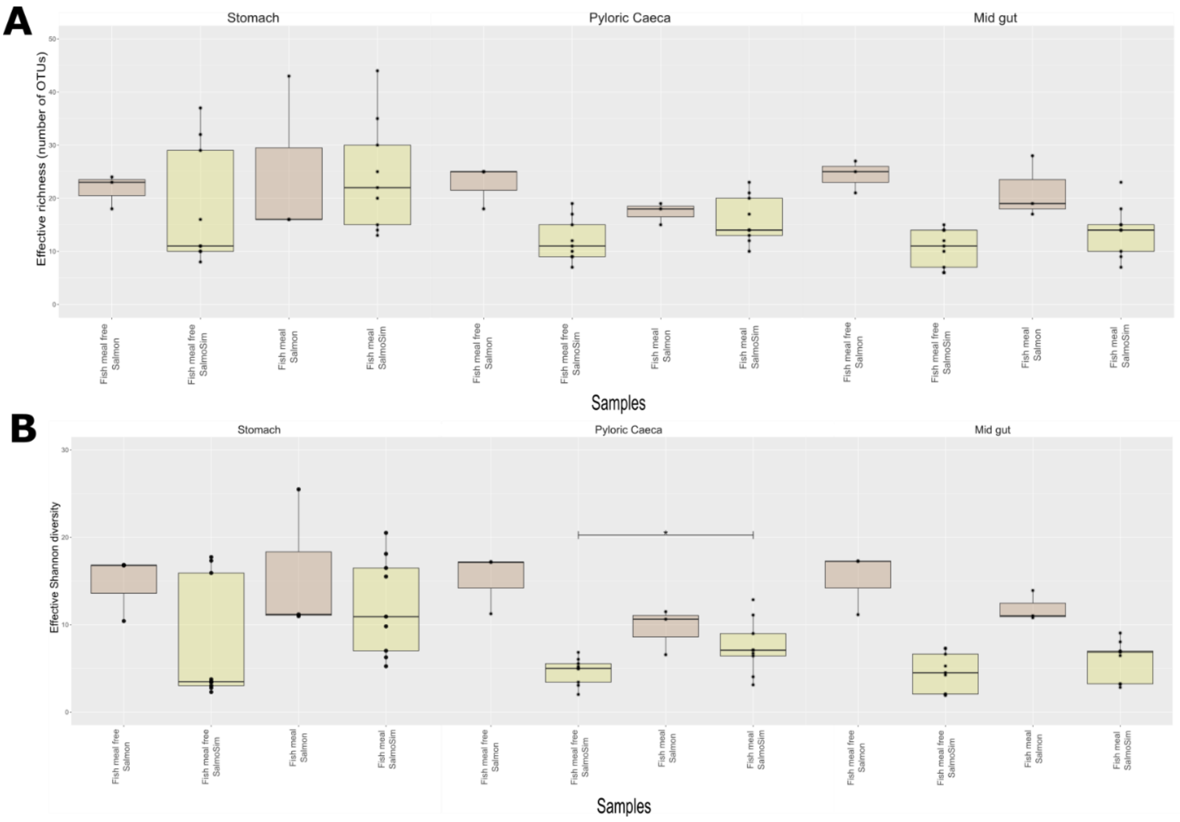


**<u>Figure 4</u> Calculated alpha-diversity metrics within different gut compartments of real salmon and SalmoSim fed on Fish meal and Fish meal free diets** *Figure above visually represents different alpha diversity outputs within different gut compartments of real salmon in red and SalmoSim in yellow (stable time points: 16, 18 and 20 fed on Fish meal, and 36, 38 and 40 fed on Fish meal free diet) fed on Fish meal and Fish meal free diets.* ***A*** *visually represents effective richness (number of OTUs),* ***B*** *represents effective Shannon diversity. The lines above bar plots represent statistically significant differences after feed change. The stars flag the levels of significance: one star (*) for p-values between 0.05 and 0.01, two stars (**) for p-values between 0.01 and 0.001, and three stars (***) for p-values below 0.001.*
