## Supplemental Table 1 for "Development of a three-compartment *in vitro* simulator of the Atlantic Salmon GI tract and associated microbial communities: SalmoSim"

**Table 1 Fish meal and Fish meal free diets composition.** *Table summarises Fish meal and Fish meal free diets composition in percentage of the feed.*

| Ingredient (% of the feed) | Fish meal | Fish meal free |
|---|---|---|
| Fish meal | 17.50 | 0.00 |
| Soya protein concentrate | 12.00 | 27.80 |
| Corn gluten | 7.00 | 7.35 |
| Wheat gluten | 10.00 | 14.34 |
| Sunflower expeller | 3.41 | 0.00 |
| Wheat | 4.81 | 11.22 |
| Beans dehulled | 10.00 | 0.00 |
| Fish oil | 15.68 | 16.99 |
| Rapeseed oil | 11.78 | 11.79 |
| Linseed oil | 3.05 | 3.20 |
| Mannooligosaccharide | 0.40 | 0.40 |
| Astaxanthin | 0.04 | 0.04 |
| Crystalline amino acids | 1.35 | 1.99 |
| Mineral premixes | 1.83 | 2.66 |
| Vitamin premixes | 0.60 | 0.73 |
