## Supplemental Table 2 for "Development of a three-compartment *in vitro* simulator of the Atlantic Salmon GI tract and associated microbial communities: SalmoSim"

**<u>Table 2</u> 16S rRNA gene-targeted group-specific primers.** *Table lists primer sets that already published and validated in the literature. All primers were used on mouse faeces samples apart from Alphaproteobacterial specific primers that were used on marine biofilm samples.*

| Group | Primer sequence | Source |
|---|---|---|
| **Bacteroidetes F** | GTTTAATTCGATGATACGCGAG | (Yang et al., 2015) |
| **Bacteroidetes R** | TTAASCCGACACCTCACGG | |
| **Firmicutes F** | GGAGYATGTGGTTTAATTCGAAGCA | |
| **Firmicutes R** | AGCTGACGACAACCATGCAC | |
| **Actinobacteria F** | TGTAGCGGTGGAATGCGC | |
| **Actinobacteria R** | AATTAAGCCACATGCTCCGCT | |
| **Gammaproteobacteria F** | TCGTCAGCTCGTGTYGTGA | |
| **Gammaproteobacteria R** | CGTAAGGGCCATGATG | |
| **Betaproteobacteria F** | AACGCGAAAAACCTTACCTACC | |
| **Betaproteobacteria R** | TGCCCTTTCGTAGCAACTAGTG | |
| **Alphaproteobacteria F** | CIAGTGTAGAGGTGAAATT | (Bacchetti De Gregoris et al., 2011) |
| **Alphaproteobacteria R** | CCCCGTCAATTCCTTTGAGTT | |
| **General Bacteria F** | ACTCCTACGGGAGGCAGCAGT | (Clifford et al., 2012) |
| **General Bacteria R** | TATTACCGCGGCTGCTGGC | |
| **Mycoplasma F** | AGCAGCCGCGGTAATACATAG | Generated by DECIPHER software* |
| **Mycoplasma R** | GAGCATACTACTCAGGCGGAT | |
| **Lactobacillus F** | CAGCAGTAGGGAATCTTCCACAA | |
| **Lactobacillus R** | CATGGAGTTCCACTCTCCTCTT | |
