## Supplemental Table 3 for "Development of a three-compartment *in vitro* simulator of the Atlantic Salmon GI tract and associated microbial communities: SalmoSim"

**Table 3** First round PCR primers used for the first round of NGS library preparation

| Primer | Name | Illumina 5′ sequencing primer (CS1/CS2) | Internal index | Heterogeneity spacer | 16S rRNA gene v1 primer |
|---|---|---|---|---|---|
| **Forward** | 27F | ACACTCTTTCCCTACACGACGCTCTTCCGATCT | Index (8bp) | heterogeneity spacer (5/3 bp) | AGAGTTTGATCMTGGCTCAG |
| **Reverse** | 338R | GTGACTGGAGTTCAGACGTGTGCTCTTCCGATCT | Index (8bp) | None | GCTGCCTCCCGTAGGAGT |
