## Supplemental Table 4 for "Development of a three-compartment *in vitro* simulator of the Atlantic Salmon GI tract and associated microbial communities: SalmoSim"

**Table 4** **Second round PCR primers used for the first round of NGS library preparation**

| Primer name | Illumina MiSeq 3′ flow cell linker (i5/i7) | External index | Illumina 5′ sequencing primer (CS1/CS2) |
| --- | --- | --- | --- |
| **Forward** | AATGATACGGCGACCACCGAGATCTACAC | Index (8bp) | CACTCTTTCCCTACACGACGCT |
| **Reverse** | CAAGCAGAAGACGGCATACGAGAT | Index (8bp) | GTGACTGGAGTTCAGACGTGTGCTC |
