## Supplemental Table 5 for "Development of a three-compartment *in vitro* simulator of the Atlantic Salmon GI tract and associated microbial communities: SalmoSim"

**<u>Table 5</u> Estimated marginal means output for a mixed-effect linear model run with different qPCR, ammonia and protein concentration values identifying differences between different gut compartments** *The table summarises the Estimated Marginal Means output for each mixed-effect linear model (Model 2) run with different values identifying the difference between compartments. A small p-value (≤ 0.05) indicates that the values between compared gut compartments are statistically different (in blue), and p>0.05 indicates that value measured is statistically similar between compared gut compartments within SalmoSim system (in green).* ***S*** *in the table refers to stomach,* ***PC*** *to the pyloric caeca, and* ***MG*** *to the midgut.*

| Value | Gut compartments between which similarity was tested | | |
|---|---|---|---|
| | **S and PC** | **S and MG** | **PC and MG** |
| **Actinobacteria** | <0.001 | <0.001 | 0.029 |
| **Alphaproteobacteria** | 0.347 | 0.003 | 0.139 |
| **Bacteroidetes** | 0.134 | <0.001 | 0.116 |
| **Betaproteobacteria** | <0.001 | <0.001 | 0.740 |
| **Firmicutes** | <0.001 | <0.001 | 0.342 |
| **Gammaproteobacteria** | 0.932 | 0.808 | 0.962 |
| **Lactobacillus** | 0.985 | <0.001 | <0.001 |
| **Mycoplasma** | <0.001 | <0.001 | <0.001 |
| **Ammonia** | <0.001 | <0.001 | <0.001 |
| **Protein** | 0.032 | <0.001 | <0.001 |
