## Supplemental Table 6 for "Development of a three-compartment *in vitro* simulator of the Atlantic Salmon GI tract and associated microbial communities: SalmoSim"

**Table 6 Beta diversity and differential abundance values from the comparison of microbial composition within different SalmoSim compartments.** *The table above summarises different beta-diversity analysis outputs calculated by using different distances: phylogenetic (unweightedbalanced and weighted UniFrac) and ecological (Bray-Curtis and Jaccards), between different SalmoSim (**A**) and real salmon (**B**) compartments: stomach (**S**), pyloric caeca (**PC**) and midgut (**MG**). Numbers represent p-values, with p-values <0.05 identifying statistically significant differences between compared groups. Equal number of samples (n=18) for each dataset were selected: real salmon (samples from the 3 gut compartments, from 3 biological replicates, for each of the 2 feeds) and SalmoSim (samples from each of the 3 gut compartments, for 3 biological replicate runs, for each of the 2 feeds (time point 20 for Fish meal and time point 40 for Fish meal free diet)).*

**A**

| Test | | S vs PC | S vs MG | PC vs MG |
|---|---|---|---|---|
| UniFrac | Unweighted (0%) | 0.145 | 0.26 | 0.843 |
| | Balanced (50%) | 0.123 | 0.185 | 0.891 |
| | Weighted (100%) | 0.112 | 0.159 | 0.936 |
| Bray-Curtis | | 0.356 | 0.298 | 0.926 |
| Jaccards | | 0.367 | 0.303 | 0.906 |

**B**

| Test | | S vs PC | S vs MG | PC vs MG |
|---|---|---|---|---|
| UniFrac | Unweighted (0%) | 0.882 | 0.742 | 0.751 |
| | Balanced (50%) | 0.909 | 0.765 | 0.85 |
| | Weighted (100%) | 0.818 | 0.735 | 0.875 |
| Bray-Curtis | | 0.767 | 0.788 | 0.734 |
| Jaccards | | 0.886 | 0.772 | 0.798 |
